# A posture subspace in primary motor cortex

**DOI:** 10.1101/2024.08.12.607361

**Authors:** Patrick J. Marino, Lindsay Bahureksa, Carmen Fernández Fisac, Emily R. Oby, Adam L. Smoulder, Asma Motiwala, Alan D. Degenhart, Erinn M. Grigsby, Wilsaan M. Joiner, Steven M. Chase, Byron M. Yu, Aaron P. Batista

## Abstract

To generate movements, the brain must combine information about movement goal and body posture. Motor cortex (M1) is a key node for the convergence of these information streams. How are posture and goal information organized within M1’s activity to permit the flexible generation of movement commands? To answer this question, we recorded M1 activity while monkeys performed a variety of tasks with the forearm in a range of postures. We found that posture- and goal-related components of neural population activity were separable and resided in nearly orthogonal subspaces. The posture subspace was stable across tasks. Within each task, neural trajectories for each goal had similar shapes across postures. Our results reveal a simpler organization of posture information in M1 than previously recognized. The compartmentalization of posture and goal information might allow the two to be flexibly combined in the service of our broad repertoire of actions.

## Introduction

We can effortlessly reach in a given direction from a wide variety of initial configurations, or postures, of the arm. Yet, due to the biomechanical properties of the limb, the muscle activity required to make the reach from different initial postures can differ in complicated ways^1^.

Somehow, populations of neurons in the brain can rapidly and flexibly combine information about arm posture and the desired goal of the movement to generate muscle commands in the moments before a movement is executed. How might posture and goal information be organized in neural activity to support their flexible combination during the range of actions we perform on a daily basis? Primary motor cortex (M1) is well-situated to play an important role in this process. It receives inputs about arm posture and movement goal from sensory and association areas and in turn provides the major source of descending projections to the spinal cord for the control of the arm and hand^2–4^ (**Figure 1**). Prior work has shown that changing the initial posture from which a movement is made can cause complicated changes in the activity of individual M1 neurons^5–9^. We reasoned that complex interactions between posture and goal at the single-neuron level could nevertheless be consistent with clear and simple organization of posture and goal information at the population level.

**Figure 1.**
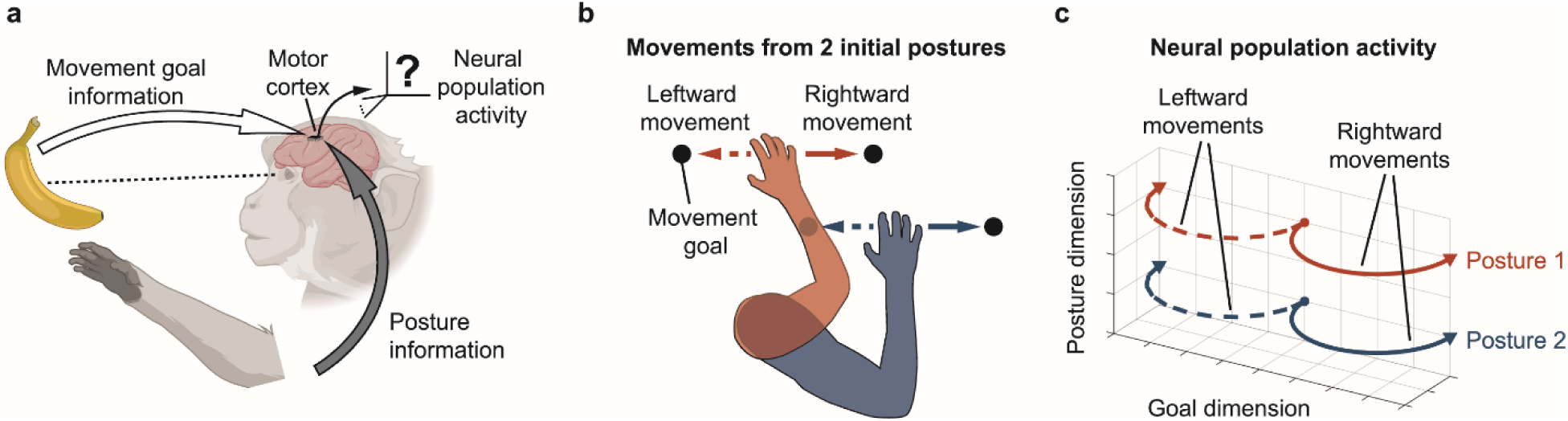
Studying how posture and goal information are organized in motor cortical activity. **(a)** M1 receives information about movement goal from upstream areas including premotor and parietal cortices^2,10^. It also receives information about arm posture from areas such as the somatosensory and parietal cortices^2,10^ and the cerebellum^4^. We sought to determine how posture and goal information were organized in M1 population activity. **(b)** We trained animals to perform tasks in which they acquired targets (i.e., movement goals) from different initial arm postures. The means of moving the cursor to the goal could be a BCI, isometric force or reaching. **(c)** We found a clear and simple population-level organization of posture and goal information: posture- and goal-related components of neural activity were separable and arranged in nearly orthogonal subspaces. Each of the four lines represents a neural trajectory corresponding to a particular arm posture and movement goal combination. Trajectory color indicates arm posture (postures illustrated in panel b), and line style (solid/dashed) indicates movement direction. Changes in posture caused a shift in neural activity that was consistent across movement directions. This shift was nearly orthogonal to the neural dimensions modulated by movement goal. Furthermore, trajectories for each movement direction had similar shapes across postures.

To investigate the structure of posture and goal information in M1 population activity, we employed a battery of behavioral tasks in which animals acquired visually-cued targets (i.e., movement goals) from a variety of initial arm postures. We used tasks that spanned a range of movement requirements. First, we examined M1 activity during a brain-computer interface (BCI) task, during which M1 is active, but in the absence of arm movements. Next, we used an isometric force task, during which muscles contract, but the arm’s posture does not change.

Finally, we examined an overt reaching task, which involves both muscular contractions and arm movements.

Across tasks, we found a clear and simple organization of posture information in M1. First, neural population activity could be decomposed into posture- and goal-related components that were separated into nearly orthogonal subspaces. Second, a single subspace could be used to decode arm posture across tasks, even though posture had different implications for each task. Third, within each task, neural trajectories for a given goal had similar shapes across postures.

Together, these results indicate the existence of a ‘posture subspace’ in M1 activity. That is, posture produces similar effects in neural population activity across a wide range of behaviors, and those effects modulate neural activity in a subspace separate from the one modulated by goal signals. The organization of posture and goal information into separate subspaces may facilitate the brain’s ability to flexibly combine the two types of information, allowing for posture information to be integrated with movement goals differently to support a wide variety of behaviors.

## Results

### Posture and goal modulate separate neural dimensions

We begin by describing how arm posture and movement goal affect neural population activity in the primary motor cortex (M1) during a BCI task. The BCI task provides a powerful tool for studying the effects of arm posture on neural activity, because during the use of a BCI, the arm is still while M1 neurons are active as they control a computer cursor directly. This allows us to examine the effects of posture and movement goal simultaneously, while minimizing the confounding effects of time-varying proprioceptive feedback generated during arm movements.

In the BCI task, animals volitionally modulated M1 activity to move a computer cursor from the center of the workspace to one of eight radially-arranged targets (i.e., movement goals) (**Figure 2a**). We used a decoder to map neural activity to cursor motion (Monkey E: position, Monkeys N and R: velocity, see **Methods**). The posture of the arm contralateral to the recording array was changed across blocks of trials by rotating the forearm about the shoulder in the transverse plane (i.e., rotating about a vertical axis through the shoulder).

**Figure 2.**
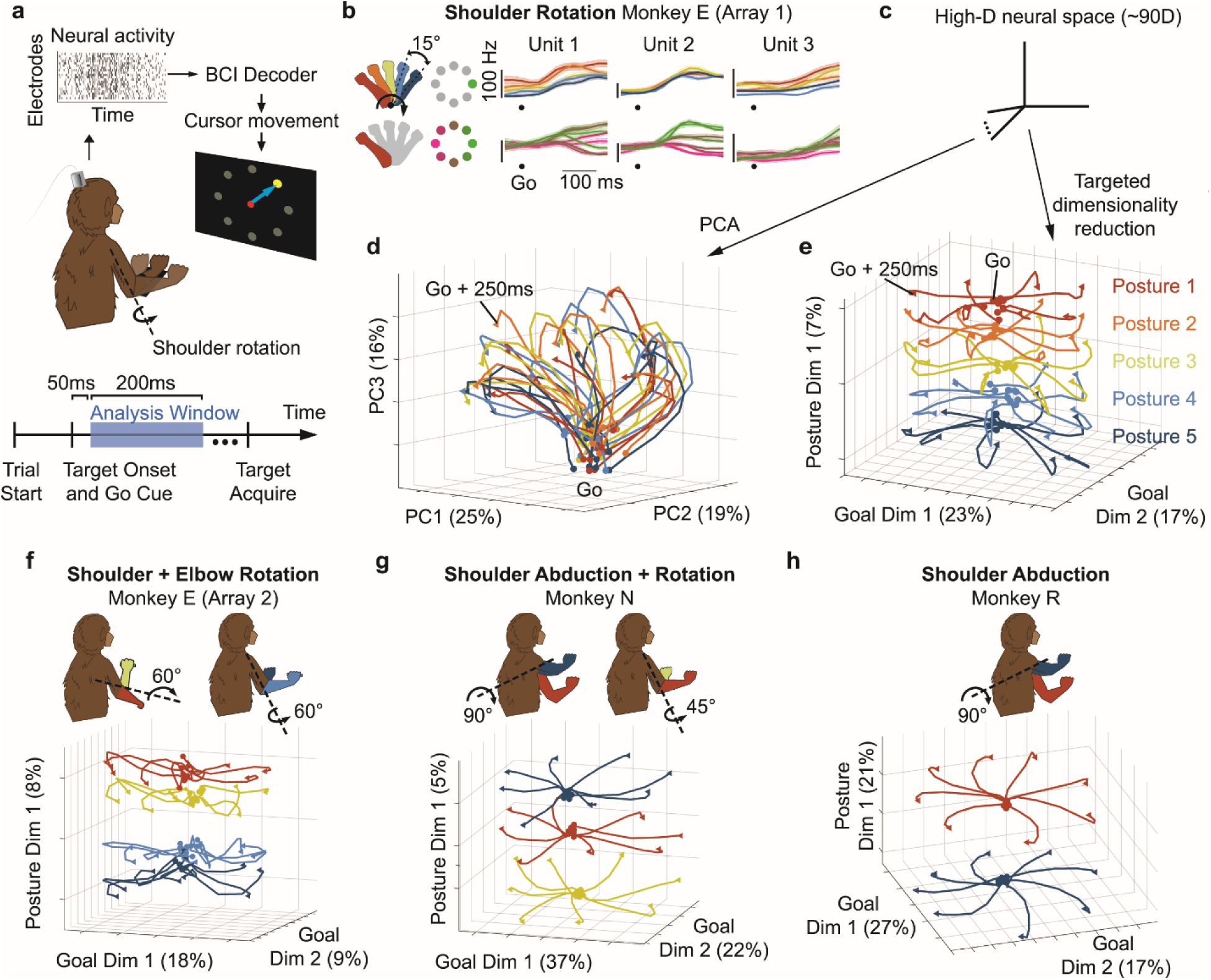
Posture and goal signals are visibly separable in neural population activity. (a) Schematic of the multi-posture BCI task. Animals volitionally modulated M1 activity to drive a computer cursor to one of eight radially-arranged targets. On each trial, a center target was acquired. Next, the center target disappeared, and one of eight peripheral targets appeared (‘target onset and go cue’). Finally, the animal acquired the peripheral target with the cursor. Neural activity from 50ms to 250ms after go cue/target onset was analyzed. The posture of the arm was changed in blocks of trials by rotating the forearm about the shoulder in the transverse plane in 15-degree increments. **(b)** (top) Trial- averaged single unit responses for the rightward BCI target for each arm posture (Monkey E, Array 1). Colors correspond to arm postures. Dot indicates target onset/go cue. Shaded error bars are +/-SEM. (bottom) Responses for all BCI target directions when the arm was in the innermost posture. Colors correspond to BCI target directions. Unit 1 is tuned to both target and posture, unit 2 is tuned primarily to target, and unit 3 is tuned primarily to posture. Note that posture tuning is visible before the go cue in units 1 and 3. **(c)** Dimensionality reduction provides different low-dimensional views of the same high- dimensional neural activity. **(d)** When visualized in the top 3 principal components, these trajectories displayed little discernible organization. Trial-averaged neural trajectories are shown for all targets and postures for Monkey E in space identified by PCA. Trajectories are colored by arm posture. Percentages along axes indicate variance of trial-averaged trajectories explained by each PCA dimension. **(e)** Here, neural trajectories are projected into the space identified by targeted dimensionality reduction (see Methods). When visualized in these dimensions, neural trajectories exhibited clear spatial structure, with dissociable effects of posture and goal. Trajectories for different targets within each posture emanated outward in different directions along ‘goal dimensions’. When the arm’s posture was changed, the shape of these neural trajectories was conserved, with all trajectories offset along ‘posture dimensions’. **(f-h)** Same format as (e), but for other monkeys and postural manipulations. (f) 60 degree elbow and shoulder rotations (Monkey E). (g) 90 degree shoulder abduction and 45 degree shoulder rotation (Monkey N). (h) 90 degree shoulder abduction (Monkey R). For all animals and postural manipulations we considered, neural trajectories exhibited similar spatial structure as in (e).

We first analyzed the responses of individual neural units during the task (**Figure 2b**). Most units exhibited mixed tuning to posture and target (e.g. unit 1), while some were tuned primarily to target (unit 2) or posture (unit 3). Postural tuning was present both before and during cursor movement, as can be seen in the responses for units 1 and 3. Many units also exhibited changes in target tuning across postures, in agreement with previously reported results for overt movement tasks^5–9^ (**Figure S1**).

We next asked how posture and goal affected the time course of neural activity across the entire population of recorded neural units. This time course can be thought of as a ‘neural trajectory’ in a high-dimensional space, where each axis of the space describes the activity of a single neural unit^11–13^. To summarize this trajectory, we projected it into a low-dimensional space identified by principal component analysis (PCA) (**Figure 2c**). We examined neural activity from 50ms to 250ms after go cue (go cue coincided with the appearance of the peripheral target that the animal acquired during the task). This window was chosen to capture goal effects in neural activity while excluding corrective cursor movements made later in the trial. When neural activity from all targets in all postures were visualized in the dimensions of greatest variance (i.e., top principal components), the neural trajectories displayed little discernible organization (**Figure 2d**).

To attempt to isolate the effects of posture and goal, we employed a targeted dimensionality reduction (TDR) approach to identify dimensions that explained posture- and goal-related variance (see **Methods**). When projected into these dimensions, neural trajectories exhibited clear spatial structure, with dissociable effects of posture and goal (**Figure 2e**). Trajectories for different targets within each posture emanated outward in different directions along ‘goal dimensions’. When the arm’s posture was changed, the shape of these neural trajectories was conserved, with all trajectories offset along ‘posture dimensions’. This created a geometry in which all trajectories from each posture were cleanly separated from those from the other postures. We asked whether similar geometry was produced by other types of arm joint rotation, including elbow rotation (Monkey E, **Figure 2f**) and shoulder abduction (Monkey N, **Figure 2g**; Monkey R, **Figure 2h**). In all cases, we found that changing the posture of the arm shifted neural trajectories along posture dimensions, while the shapes of the neural trajectories for each target were conserved.

Before concluding that posture and goal modulate separate neural dimensions, we considered several alternative explanations for the observed geometry in neural trajectories. The geometry was not a byproduct of TDR, although TDR made it more apparent; it was also visible for neural trajectories from the same target direction in the top principal components (**Figure S2**).

Changes in BCI decoders across postures did not explain the observed postural effects (**Figure S3**). Similarly, drift in neural signals over time could not account for the effects of posture (**Figure S4**). After ruling out these alternative explanations, we can conclude that posture and target have dissociable effects on neural population activity during BCI control: changes in posture impact neural trajectories by offsetting them along posture dimensions, while changes in target alter the direction in which trajectories emanate along goal dimensions.

### Posture and goal subspaces are nearly orthogonal

Visualizations of neural activity from the BCI task (**Figure 2**) suggested that posture and goal have separable effects on neural population activity in M1, and that these effects modulate different neural dimensions. Previous work has suggested that compartmentalization of sensory and motor information into orthogonal subspaces could support flexible sensorimotor integration^14,15^. Separating neural activity across postures may allow different dynamical flow fields of neural activity to be learned in each posture^11,16–18^. Additionally, compartmentalization may prevent incoming sensory information from prematurely affecting ongoing motor commands^14,15^. To determine the extent to which this compartmentalization principle applies to posture and goal information, we asked how different the posture and goal dimensions were. At one extreme, these dimensions could be orthogonal (**Figure 3a, top**). At the other extreme, they could be aligned (**Figure 3a, bottom**).

**Figure 3.**
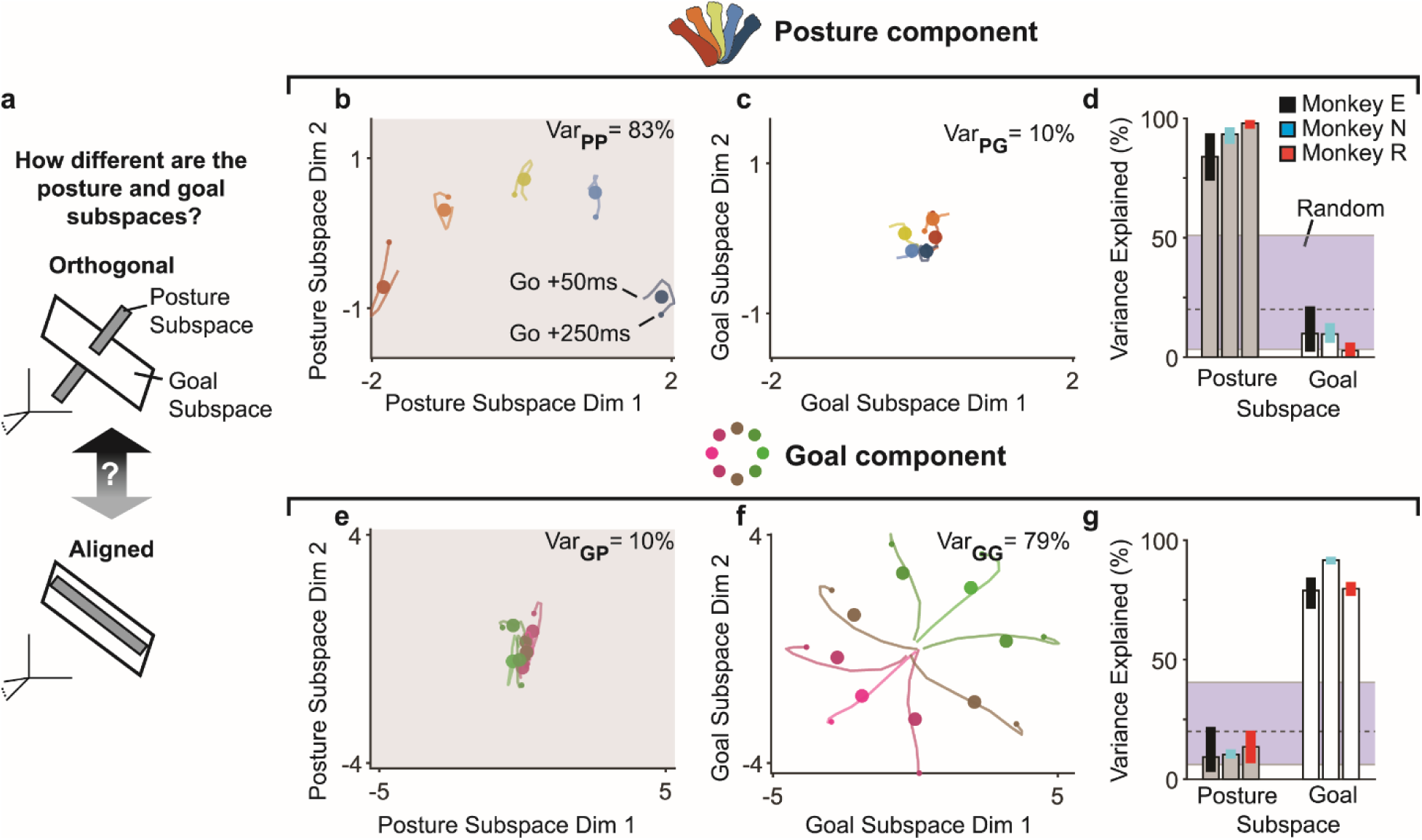
Posture and goal subspaces are nearly orthogonal. (a) How similar are the posture and goal subspaces? At one extreme, they may be orthogonal (top) while at the other, they may be aligned (bottom). In this schematic, the posture subspace is shown as 1- dimensional to illustrate orthogonality and alignment. (**b**) Projection of the posture component of neural population activity into the 2-dimensional posture subspace for the same session as shown in Figure 2e. Each trace is the marginalization over target directions for one posture (see Methods). Traces are colored by posture. Small dots indicate the end of the trajectory, and large dots represent the mean across time. These means are only shown for visual clarity; quantifications were performed on the marginalized time courses. For this session, the posture subspace captured 83% of the posture-related variance. **(c)** Projection of the posture component into the 2-dimensional goal subspace. The goal subspace captured only 10% of the posture-related variance for this session. **(d)** Posture-related variance captured by posture and goal subspaces for each animal. Data from all sessions and postural manipulations were combined for each monkey (Monkey E: four shoulder rotation sessions and one session in which both the shoulder and elbow were rotated; Monkey N: two sessions in which the shoulder was separately rotated and abducted; Monkey R: one session of shoulder rotation and one session of shoulder abduction; see Methods). Bar heights are means across sessions and error bars indicate the 2.5^th^ to 97.5^th^ percentile of the bootstrapped sampling distribution. The gray dashed line and shaded area indicate the mean and 2.5^th^ to 97.5^th^ percentile of posture-related variance captured by randomly drawn subspaces (see Methods). For each animal, the goal subspace captured much less posture-related variance than the posture subspace (p < 10^-24^, bootstrap test), and the amount of posture-related variance captured by the goal subspace was on the low end of the random subspace distribution. Together, these indicate that the posture and goal subspaces are nearly orthogonal. (**e-g**) Same format as (**b-d**), but for goal-related variance. Across animals, the posture subspace captured much less goal-related variance than the goal subspace (p < 10^-27^, bootstrap test), providing further evidence that the posture and goal subspaces are nearly orthogonal.

To quantify the alignment of the posture and goal dimensions, we performed a cross-projection alignment test^19^. We first decomposed trial-averaged neural activity into a posture component (by averaging neural trajectories over different targets; see **Methods**), and goal component (by averaging neural trajectories over different postures). We then identified the ‘posture subspace,’ defined as the dimensions that captured the greatest posture-related variance, by applying PCA to the posture component (see **Methods**). We separately identified the ‘goal subspace’ by applying PCA to the goal component. We next projected the posture component into each subspace and measured the amount of variance captured in a cross-validated manner. If the two subspaces were perfectly aligned, they would capture the same amount of posture-related variance. If they were orthogonal, then the goal subspace would capture little posture-related variance. We then repeated this procedure with the goal component.

We found that the posture subspace captured the majority of posture-related variance, whereas the goal subspace captured very little posture-related variance. This is an indication that the posture and goal subspaces were nearly orthogonal. For example, in the session shown (Monkey E), a 2-dimensional posture subspace captured 83% of the posture variance (**Figure 3b**), whereas a 2-dimensional goal subspace captured only 10% of the posture variance (**Figure 3c**). This result was consistent in all sessions for all animals (**Figure 3d**). In fact, the amount of posture variance captured by the goal subspace was on the low end of the amount captured by randomly-drawn subspaces. Similarly, we found that the posture subspace captured little goal-related variance. In the session shown, the 2-D posture subspace captures 10% of the goal variance (**Figure 3e**), whereas the 2-D goal subspace captures 79% of the goal variance (**Figure 3f**). This effect was also observed in all sessions for all animals (**Figure 3g**).

When we manipulated multiple arm joints within the same session, the posture and goal subspaces remained orthogonal, and the posture subspaces for each joint were orthogonal to one another (**Figure S5**).

These results indicate that, during BCI control, neural population activity can be decomposed into posture- and goal-components, and that these components modulate nearly orthogonal subspaces of neural activity. We next sought to determine whether these findings extended to overt tasks requiring muscular contraction.

### Posture and goal also modulate separate neural dimensions during overt tasks

The BCI paradigm enabled us to isolate the effects of posture and goal on neural activity, because during BCI control, M1 is active though the arm does not move, and the muscles need not contract. This removes many of the confounding factors present during overt movement, such as time-varying proprioceptive feedback and movement commands that may depend on initial arm posture. So far we have shown that posture and goal modulate distinct subspaces of M1 population activity, at least during BCI control. Does this finding generalize to ‘overt’ behaviors (i.e., behaviors requiring muscular contraction)? We next asked whether a posture subspace was present during overt isometric force and reaching tasks.

In the isometric force task, muscle activation is required for task success, but limb movement is restricted, which minimizes the influence of time-varying proprioceptive feedback (**Figure 4a**). The animal exerted upward or downward forces on a handle attached to a force transducer to move a computer cursor to a target displayed on the screen. Between blocks, arm posture was changed by rotating the forearm about the shoulder in the transverse plane. We analyzed neural activity from 50ms to 250ms after go cue to include strong target effects on neural activity while excluding as much as possible time-varying proprioceptive feedback from the contracting muscles (see **Methods**).

**Figure 4.**
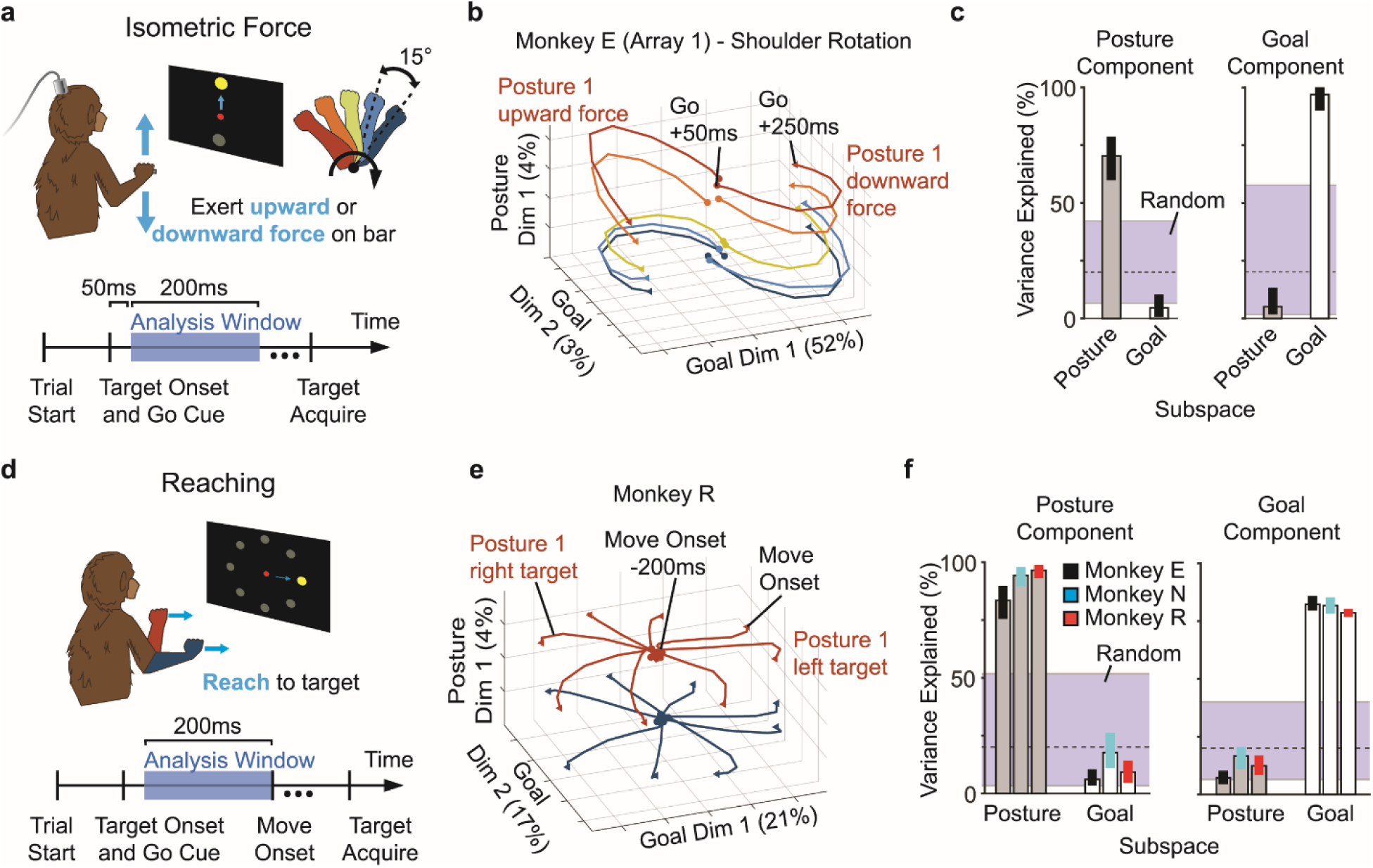
Posture and goal also modulate separate neural dimensions during overt tasks. (a) Schematic of the isometric force task. Monkey E (Array 1) exerted upward or downward forces on a handle that was fixed to a force transducer. Forces recorded by the transducer were mapped to cursor position. On each trial, a central target was first acquired. Next, the center target disappeared, and an upward or downward peripheral target appeared (‘target onset and go cue’). The animal exerted forces to drive the cursor to acquire the peripheral target (4.75N of upward or downward force corresponded to the upward or downward target center, respectively). Neural activity from 50ms to 250ms after go cue/target onset was analyzed. The posture of the arm was changed across blocks of trials by rotating the forearm about the shoulder in the transverse plane in 15-degree increments. **(b)** Trial-averaged neural trajectories for all targets and postures for the isometric force task, plotted in the space identified by TDR (see Methods). Neural trajectories exhibited clear spatial structure, with dissociable effects of posture and goal. Trajectories are colored by arm posture. Percentages along axes indicate variance of trial-averaged trajectories explained by these dimensions. **(c)** Posture- and goal-related variance captured by the posture and goal subspaces for the isometric force task. Same format as Figure 3d (left) and Figure 3g (right). As with the BCI task, the goal subspace captured much less posture-related variance than the posture subspace (left, p < 10^-20^, bootstrap test), and posture subspace captured less goal-related variance than the goal subspace (right, p < 10^-20^, bootstrap test). Together, these support the notion that the posture and goal subspaces are nearly orthogonal. **(d)** Schematic of the reaching task. On each trial, a center target was acquired. Next, the center target disappeared, and one of eight radially-arranged peripheral targets appeared (‘target onset and go cue’). Finally, the animal made a reach to acquire the peripheral target. We analyzed neural activity in the 200ms before movement onset. Posture was changed in randomized blocks of trials by adjusting the initial location of the hand in the frontoparallel plane. Visual feedback was matched across postures. Task design for Monkey R is shown; task design varied slightly for other animals (see Methods). **(e)** Trial-averaged neural trajectories for all movement goals and postures for one example session of the reaching task, plotted in the space identified by TDR. **(f)** Posture- and goal-related variance captured by the posture and goal subspaces for the reaching task. Same format as Figure 3d (left) and Figure 3g (right). Again, the goal subspace captured much less posture-related variance than the posture subspace (left, p < 10^-52^, bootstrap test), and posture subspace captured less goal-related variance than the goal subspace (right, p < 10^-63^, bootstrap test), indicating that the posture and goal subspaces are nearly orthogonal.

When we examined neural population activity in the isometric task, we again observed that neural trajectories for each target had similar shapes across postures and were separated along posture dimensions (**Figure 4b**). We repeated the subspace alignment analysis and again found that posture and goal subspaces were nearly orthogonal (**Figure 4c**). These observations indicate that a posture subspace is also evident during an isometric task, as it was for BCI control.

We next asked whether posture and goal information were also separated during a center-out reaching task performed from different initial arm postures. As similar tasks have been shown to produce substantial changes in muscle activation and neural activity across postures^5^, we wondered if the clean separation of posture and goal effects would break down during an overt reaching task, or if our ability to see it would be obscured. In this task, animals made reaches to one of eight radially arranged targets. The initial posture of the arm was changed across blocks of trials by changing the initial location of the hand in the frontoparallel plane (**Figure 4d**).

Reaches were matched in direction and length from each initial hand position. Visual feedback was matched across postures so that the instructed initial hand position always corresponded to the center of the screen. We analyzed neural activity in the 200ms preceding movement onset to exclude time-varying proprioceptive feedback.

Even during overt reaching, we observed clear organization of posture and goal information in neural population activity (**Figure 4e, Figure S6**). Posture and goal subspaces were nearly orthogonal (**Figure 4f**). We again noticed that neural trajectories for a given target seemed to have a similar shape (e.g., **Figure 4e**, red and blue trajectories for the rightward target). We return to this point below for further quantification. We note that once the arm begins moving, activity in the posture subspace does not closely track the position of the hand (**Figure S7**), possibly due to muscle lengths changing in complicated ways during reaching.

Our consistent results across tasks suggest that the separation of posture and goal information into distinct subspaces is an organizing principle of motor cortex. It is easiest to see in the BCI case, but also present during behaviors that engage the muscles and move the arm.

### The posture subspace is shared across tasks

Across a variety of tasks, we found that posture and goal information were separated into distinct neural subspaces. We next asked how the posture subspaces from different tasks were related. It could be the case that there is a separate posture subspace for each of our tasks.

This is plausible because, in each task, the initial arm posture has different implications for the upcoming muscle commands. Alternatively, it could be that there is a posture subspace that is common to all tasks. This might be the case if the posture subspace purely reflects joint-based proprioceptive inputs to M1. To directly compare the encoding of posture across tasks, we trained Monkey E to perform a ‘multiple tasks’ paradigm, in which all three of the tasks were performed within a single experimental session (**Figure 5a**).

**Figure 5.**
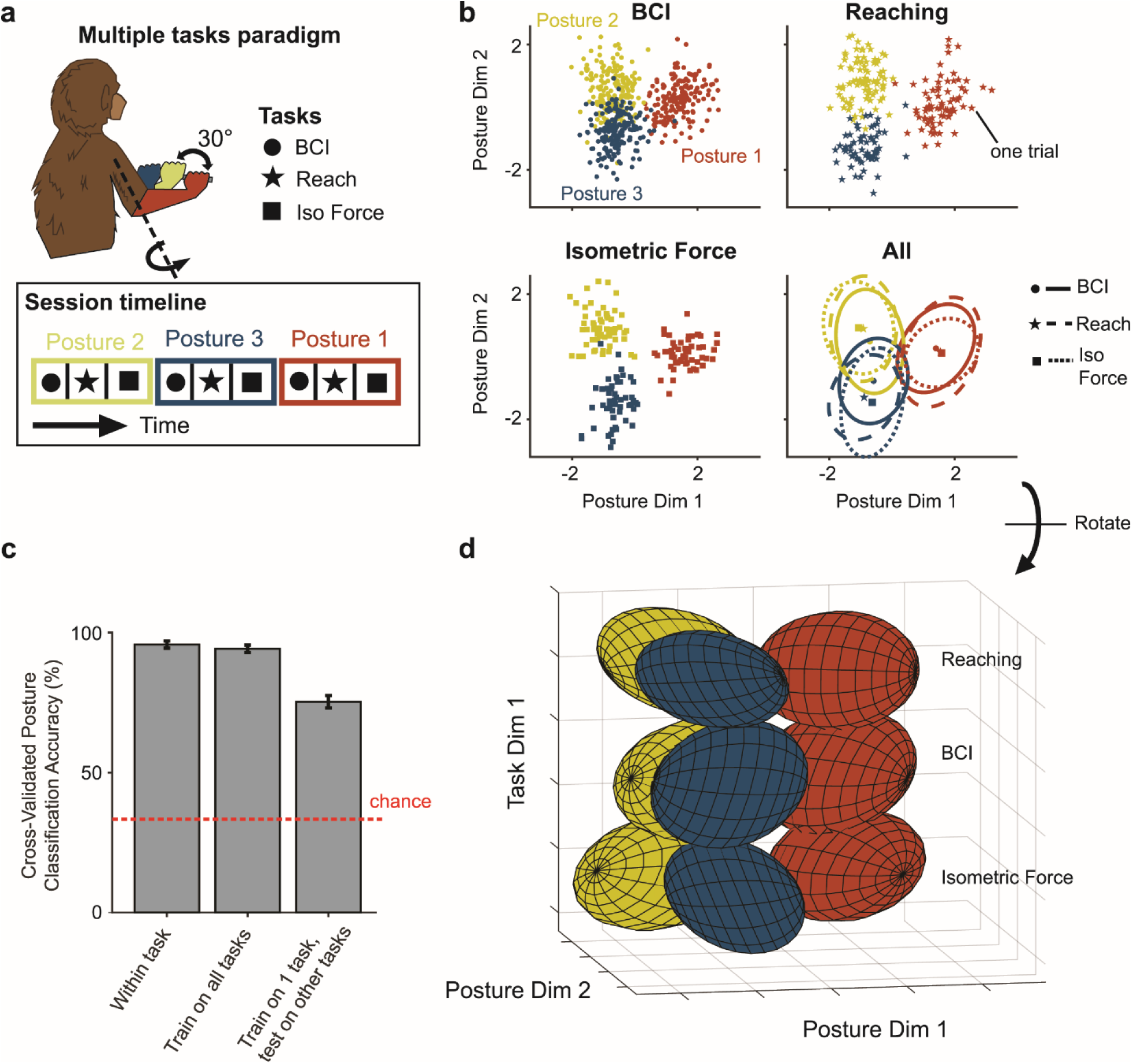
The posture subspace is shared across tasks. (a) In the multiple tasks paradigm, Monkey E (Array 1) performed the BCI task, the isometric force task, and the reaching task in blocks of trials on the same day. The tasks were modified so that the same three initial postures were matched across tasks (see Methods). The posture of the arm was changed in blocks of trials by rotating the forearm about the shoulder in the transverse plane in 30-degree increments. Blocks of each task were performed in each posture before changing postures. **(b)** Projections of mean neural activity for individual trials (one point per trial) onto the posture subspace identified by linear discriminant analysis (LDA) on data from all tasks. The first three panels each show neural activity from one task. The lower-right panel shows neural activity from all tasks together, with ellipses covering 95% of the distribution for each task and posture. Axes are the same for each panel. Class labels for LDA were determined by arm posture, irrespective of task or target direction. Projections of neural activity for each task were overlapping in the posture subspace (lower right panel), indicating that it may be possible to classify arm posture from a single subspace, regardless of task. **(c)** Arm posture can be accurately classified from motor cortical activity across different tasks. To assess the consistency of postural signals across tasks, we calculated cross-validated posture classification accuracy in three different ways (for details, see Methods). First, we trained and tested separate classifiers within each of the three tasks (*Within task*, left bar). Bar height and error bar are mean and SEM of the classification accuracies across 15 test folds (five folds for each of the three tasks). This effectively establishes an upper limit for classifier performance against which we can compare across-task classification performance. Second, we trained a single classifier on neural activity from all tasks, then tested on data from each task individually (*Train on all tasks*, middle bar). We observe almost the same classifier performance when compared with the within task classification, suggesting that we can identify a consistent posture subspace across tasks. Third, we trained classifiers on neural activity from one task and then tested on data from the other tasks (*Train on 1 task, test on other tasks*, right bar). Classifiers performed well above chance (red dashed line, 33.3%), again suggesting a consistent posture subspace across tasks. **(d)** Distributions of neural activity from each posture and task combination, visualized in a 3-dimensional space formed from the posture subspace from (b) (horizontal plane) and a ‘task dimension’ (vertical axis). Ellipsoids contain 95% of the neural activity in the distribution. To identify the task dimension, we used LDA to separate neural activity by task, ignoring arm posture and target direction. This procedure identifies two task dimensions, as there were three task labels. One dimension was chosen manually from within the space formed by these two task dimensions and combined with the two posture dimensions (see Methods). We then used QR decomposition to orthonormalize these three dimensions. The resulting visualization demonstrates that neural activity maintains a consistent projection into the posture subspace, even as activity differs strongly across tasks.

We asked whether it was possible to identify a single subspace from which arm posture could be read out across tasks. To test this, we averaged neural activity within the analysis window for each task, resulting in one observation (i.e., one spike count vector) per trial. We then used linear discriminant analysis (LDA) to identify a subspace that separated neural activity from all tasks by posture (see **Methods**). Projections of each task’s neural activity into the subspace identified by LDA were qualitatively similar (**Figure 5b**), suggesting that a single posture subspace might be common to all tasks.

To quantify this, we attempted to classify posture from neural activity using a single classifier across tasks. We used the means and covariance of the neural population activity from LDA to classify posture. To establish a baseline for classifier performance, we first trained classifiers on each task separately, and found that the cross-validated prediction accuracies were nearly perfect (96.7% correct; **Figure 5c**, left bar). When we used a single classifier for all tasks, there was hardly any degradation in performance (95.4% correct; **Figure 5c**, middle bar). An even stronger test of the consistency of the posture subspace across tasks is to train a posture classifier on one task and test on others. When we did this, the classifiers still performed well above chance levels, with a moderate decrement in performance (77.8% correct; **Figure 5c**, right bar).

A trivial explanation for our ability to decode posture using a single subspace across tasks is that overall neural activity is similar across tasks. However, we were able to identify other neural dimensions that strongly separated neural activity across tasks, indicating that neural activity for each task was starkly different (**Figure 5d**; see **Methods** for details on how these dimensions were identified). This suggests that the posture subspace is consistent across tasks, even though overall neural activity is different across tasks. Together, these analyses support the existence of a stable posture subspace in M1’s population activity.

### Neural trajectories in each task have similar shapes across postures

So far, we have shown that posture and goal information are separated in M1 neural population activity across a variety of tasks, and that the posture subspace is consistent across tasks.

When examining neural population activity within each task, we noticed that neural trajectories for each target seemed to have a similar overall shape, regardless of arm posture. Similarity of trajectory shape might indicate that goal-related neural motifs are reused across postures, potentially allowing for ‘composition’ of movement commands^20,21^. Therefore, we next sought to precisely quantify the extent to which trajectories changed shape across postures.

To do so, we measured the difference (as the mean Euclidean distance) between pairs of trajectories before and after applying a translation to align their means (**Figure 6a**; for complete illustration of procedure, see **Figure S8**). This translation accounts for the offset in population space between the pair of trajectories, including any offset due to the influence of posture. We compared the difference after translation to the difference before translation for each pair of trajectories (**Figure 6b**). If trajectories do not reshape, then applying this translation would remove any difference between them, yielding completely overlapping trajectories. This would result in a low ‘difference after translation,’ regardless of the ‘difference before translation.’ Alternatively, if the pair of trajectories had different shapes, the translation procedure would account for some (but not all) of the difference between them, resulting in a higher difference after translation.

**Figure 6.**
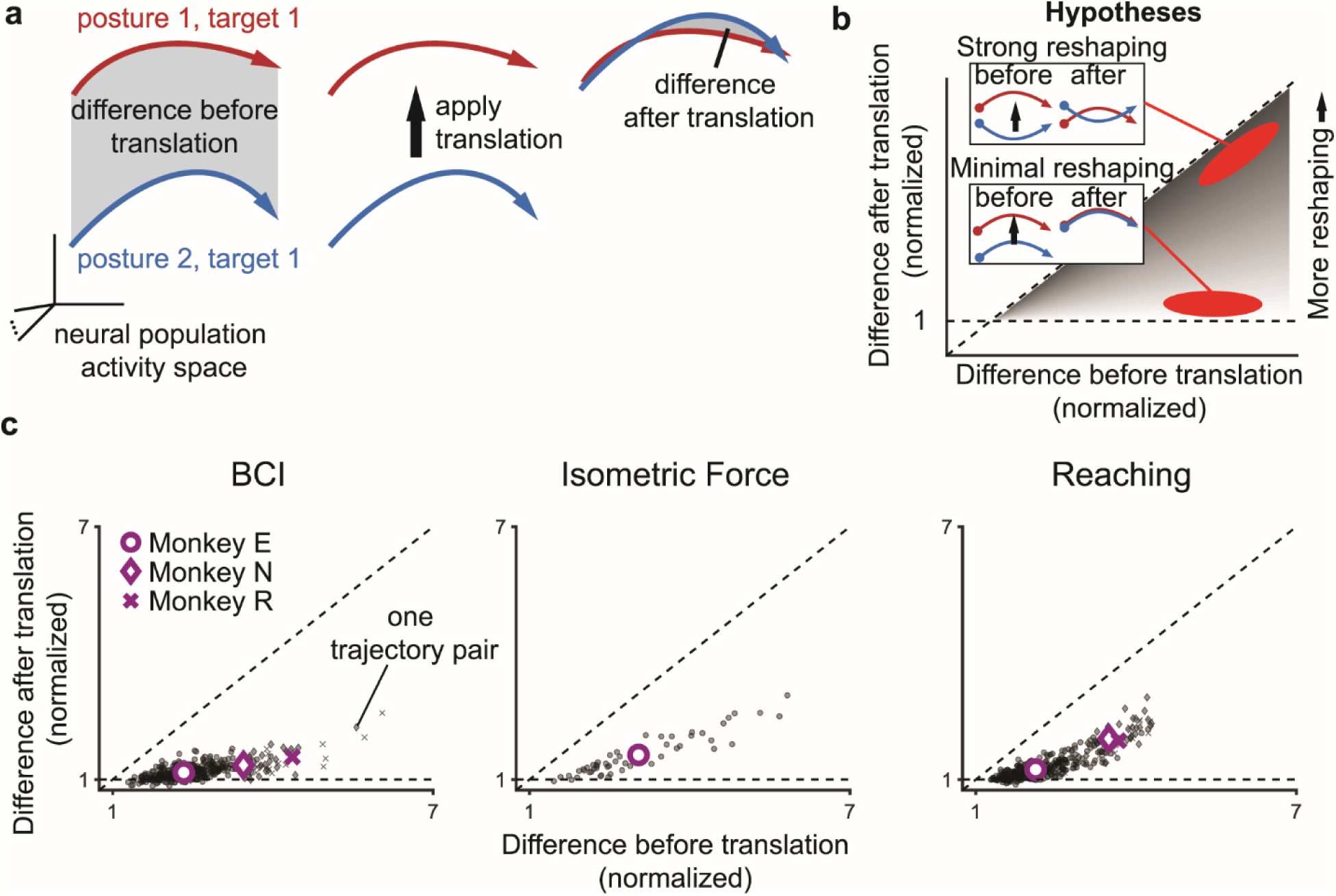
Neural trajectories in each task have similar shapes across postures. (a) Method for assessing the similarity of neural trajectories across postures (see Methods, Supplementary Figure 8). We measured the difference between pairs of trajectories before and after translating them to bring them into alignment. Our metric for the difference between a pair of trajectories was the mean of the Euclidean distances between corresponding time points on each trajectory. **(b)** Trajectories might exhibit *strong reshaping* across postures (top), in which case the difference before and after translation would be similar, producing points near the diagonal. Alternatively, trajectories might exhibit *minimal reshaping* across postures (bottom), in which case the difference after translation would be near 1, regardless of the difference before translation. **(c)** Neural trajectories for each task have similar shapes across postures. Each panel shows the differences before and after translation for all monkeys and tasks. Each small, shaded marker represents one comparison for one experimental session (e.g., posture 1, target 1 vs. posture 2, target 1). The larger markers indicate the mean of all comparisons for each monkey (circle, Monkey E; diamond, Monkey N; X, Monkey R). In the BCI task, all differences after translation are near 1, regardless of the difference before translation. This indicates minimal trajectory reshaping in the BCI task. In the isometric force and reaching tasks, differences after translation were slightly higher, although still well below the diagonal. This indicates that trajectories still had broadly similar shapes across postures, with some reshaping that might be explained by differences in muscle activity or sensory feedback across postures.

Across all tasks, we found that removing the offset between trajectories accounted for the majority of the difference between them (BCI, Monkey E: 89% reduction, Monkey N: 85%, Monkey R: 83%; Isometric Force, Monkey E: 70%; Reaching, Monkey E: 73%, Monkey N: 58%, Monkey R: 63%), resulting in low values for differences after translation (**Figure 6c**). We noticed that differences after translation were higher for isometric force and reaching than they were for the BCI task. This means that trajectories exhibit larger changes in shape across postures for these overt tasks than they do for the BCI task. This could be due to differences in time-varying proprioceptive feedback or outgoing muscle commands across postures that are present in the overt tasks, but not BCI task. These differences notwithstanding, within each task, the shapes of trajectories were largely conserved across postures, supporting the possibility that M1’s posture and goal information could serve to flexibly compose motor commands.

## Discussion

Here we report the existence of a “posture subspace” in M1. By incrementally varying arm posture in BCI, isometric force, and reaching tasks, we discovered that posture and movement goal information are mostly confined to separate subspaces of M1 population activity. The posture subspace is preserved across tasks. Finally, for any given task and movement direction, neural trajectories had similar shapes across postures. Our results reveal a simpler organization of posture information in M1 than previously recognized.

Prior work has suggested that the separation of distinct neural signals into orthogonal subspaces may be advantageous for sensorimotor integration^14,15,22^, as well as for learning^18,23,24^, motor control^19,20,25–27^, working memory^28,29^, and decision making^30–33^. The separation of posture and goal signals in M1 may provide similar benefits, allowing M1 to flexibly integrate posture information when generating movement commands. We propose three possible benefits. First, the separation of posture and goal signals could allow goal signals to be re-used across postures, thereby enabling compositional coding of movement commands and facilitating generalization to new postures^20,21^. This is consistent with our observation that neural trajectories have broadly similar shapes across postures, even for reaches occurring in different parts of the workspace. Second, the separation of posture and goal signals could enable different dynamical flow fields of neural activity to be learned in different postures^11,16–18^. We found that neural trajectories had similar overall shapes in each posture, but the shapes were not identical (**Figure 6c**). These small changes in shape could reflect slight differences in flow fields across postures. Indeed, recent work has shown that muscle-related components of M1 activity explain only a small fraction of the variance in overall population activity^34^, suggesting that slight differences in neural flow fields could be sufficient for producing the muscle activity required in each posture^35^. Third, the separation of posture and goal signals could allow posture information to enter M1 without immediately causing limb movement. This is because posture signals could avoid M1’s ‘output-potent space,’ the dimensions which cause muscle output^25^. By sequestering posture information from the output-potent space, M1 could process posture information before it affects movement^14,15^, allowing it to be used in a goal-dependent way^36^, as opposed to being mapped one-to-one to muscle activity, as can happen for a simple reflex.

How might the posture subspace arise? It is likely that the posture signal we observed originates from proprioceptive stretch receptors in the muscles^37,38^. For most of our tasks, the monkey’s arm was resting passively on a handle during the analysis window, so it is unlikely that the posture signal was an efferent signal related to active postural maintenance. Indeed, previous work has found that changes in the passive configuration of the arm influence M1 activity, even when the muscles are relaxed^37^. However, this does not explain why the posture subspace is nearly orthogonal to the goal subspace. One possible explanation for the observed near orthogonality is that proprioceptive signals arrive at M1 via different anatomical pathways (e.g., through somatosensory cortex) than goal signals (e.g., through premotor cortex). These distinct pathways could form disparate, independent patterns of synaptic connections with M1. Thus, such projections could drive population activity patterns that are orthogonal to each other. Another possibility is that orthogonal subspaces are set up through learning to exploit the computational benefits enumerated above^23^.

The population-level organization of posture and goal information that we report is simpler than previously recognized, as posture influences the activity of single units in M1 in complicated ways. For example, as reported in previous studies^5–9,37,39,40^, we observed changes in the directional tuning of individual neural units across postures, as well as heterogeneous offsets in the overall firing rate of individual units across postures (cf. **Figure S2**). Yet, these seemingly- complicated changes in activity with posture are consistent with a far simpler organization in population activity space, made evident with dimensionality reduction techniques^41,42^. We found that a single neural plane could capture goal signals across postures, and that neural population trajectories had similar shapes across postures. We proceeded to incrementally vary posture in a variety of behavioral tasks, which enabled us to discover that the postural signals are compartmentalized from goal signals in a consistent way across tasks.

Although this organization was visible across a wide range of tasks, there may be scenarios in which the organization is less apparent. For example, it may be less apparent in tasks requiring drastic changes in muscle activity across postures (e.g., Kakei et al. 1999^6^, Oby et al. 2013^7^).

This is suggested by our finding that neural trajectories reshape more across postures in the reaching task, which required changes in muscle activity across postures, than they did in the BCI task, which did not involve arm movements.

For users of clinical BCIs, it is important that postural signals do not impede control of the BCI. Postural signals can be present for BCI users who have experienced incomplete transection of spinal sensory pathways^40^, and BCI users may change posture if they retain some degree of residual motor control or if their body is adjusted by a caregiver. Our results suggest that it may be possible to design BCI decoders that are robust to posture signals. We found that posture signals are mostly confined to the posture subspace, and that goal signals are similar across postures. This implies that if enough postures are sampled to accurately estimate the goal subspace, a single decoder can be trained to decode movement intention from all of them. After training, the decoder would not need to be recalibrated each time posture changes. Another implication of our results is it may also be possible to design decoders that are robust to somatosensory signals generated by electrical stimulation, provided that the stimulation is delivered in such a way that it avoids the goal subspace.

Overall, our work adds posture to a growing list of non-motor signals that influence M1’s activity^43^, including arousal^44^, reward^45^, uncertainty^27^, somatosensory feedback^15,37,46^, visual feedback^47,48^, and memories^18,49^. Ongoing work will continue to determine how these non-motor signals influence M1’s neural population dynamics, and how this impacts decision making, learning, and ultimately, the movement that is produced.

## Acknowledgments

We thank Rob Gaunt, Neeraj Gandhi, Hillary Wehry, Sam Snyder, and Raeed Chowdhury for helpful discussions, Kevin Thiel for assistance building experimental equipment, and Jenn Sakal for help with animal care.

## Author contributions

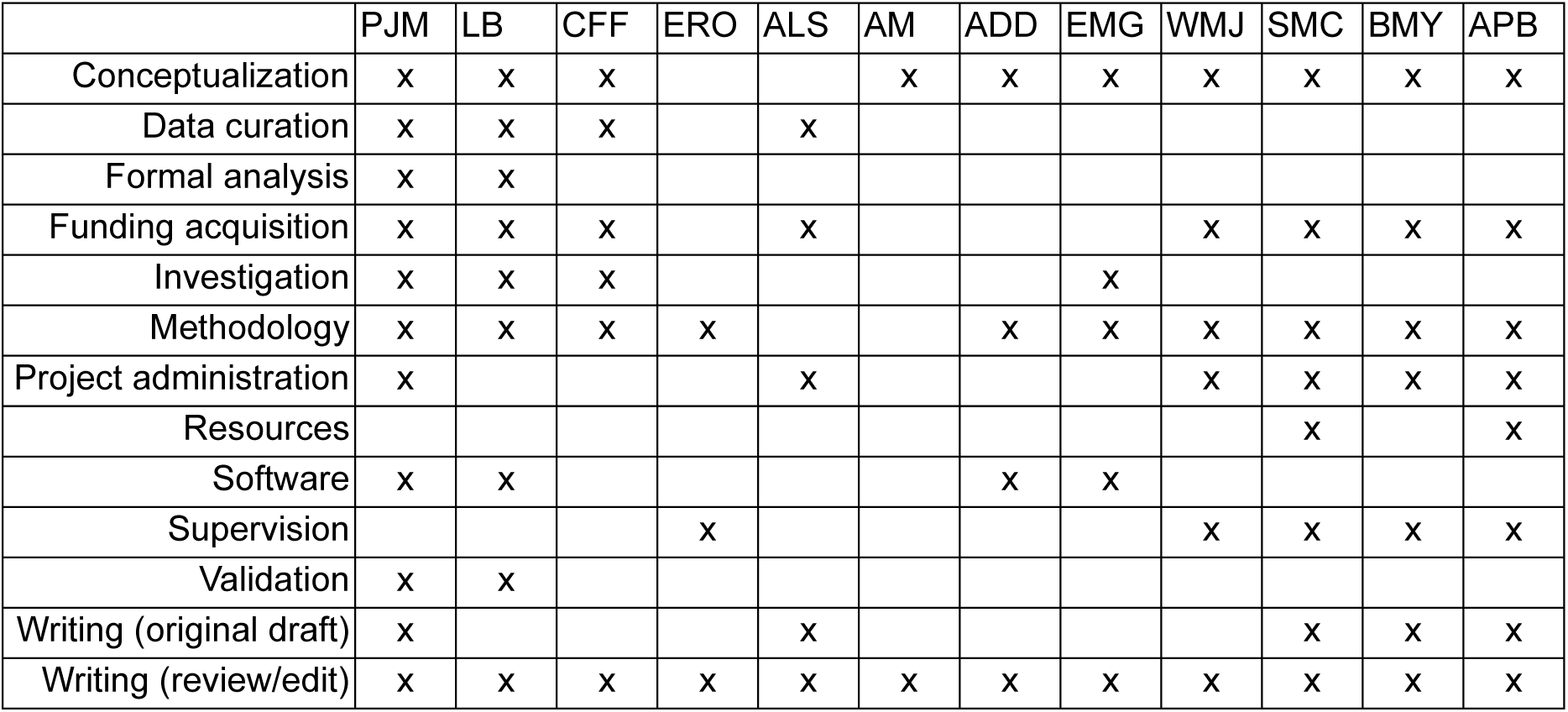

## Methods

### Resource availability

Requests for materials should be directed to and will be fulfilled by the lead contact, Aaron Batista.

### Materials availability

This study did not generate new unique reagents.

### Data and code availability

Data will be made available at the time of publication. Analysis code can be found at https://github.com/pmarino162/posture-git-repo

### Experimental subjects and details

Three adult male rhesus macaques, Monkey E (9.2kg, 10 years old), Monkey N (10.7kg, 7 years old), and Monkey R (17.1kg, 7 years old), were used in this study. The experiments for this study were conducted at the University of Pittsburgh (Monkey E) and Carnegie Mellon University (Monkeys N and R). All animal handling procedures were approved by the University of Pittsburgh Institutional Animal Care and Use Committee or the Carnegie Mellon University Institutional Animal Care and Use Committee, respectively. All data were analyzed using MATLAB 2019a.

### Neural recordings

We recorded from the proximal arm region of the motor cortex of all animals using microelectrode ‘Utah’ arrays (Blackrock Microsystems). Prior to array implant surgeries, all animals were trained to perform center-out reaching tasks. Monkey E was also trained to perform isometric force and delayed center-out reaching tasks. We implanted arrays in the hemisphere contralateral to the trained reaching arm.

Monkey E was first implanted with a 96-channel array straddling the shoulder regions of primary motor cortex (M1) and dorsal aspect of the premotor cortex (PMd) of the left hemisphere (Array 1), and a series of experiments were conducted using the right arm (see details below). Later, after training Monkey E to use his left arm in a reaching task, two separate 64-channel arrays were implanted, one in PMd and one in M1 of the right hemisphere, and second series of experiments were conducted with the left arm. We refer to these data collectively as Monkey E, Array 2. Monkey N was implanted with one 96-channel array in M1 of the left hemisphere.

Monkey R was implanted with two adjacent 96 channel arrays (arranged medio-laterally) in M1 of the left hemisphere. In this study, we make no distinction between recordings from M1 versus PMd and include all recordings for all analyses.

Microelectrode array analog voltage signals were amplified, bandpass filtered, and digitized using a multichannel acquisition processor system (Tucker Davis Technologies, Monkey E; Plexon, Monkey N; Blackrock Microsystems, Monkey R). Waveform data were converted to spiking events by thresholding at 3x (Monkey E) or 3.5x (Monkeys N and R) the root mean square voltage for each channel. We did not include data from an electrode if the threshold crossing waveforms did not resemble action potentials or if the electrode appeared to be electrically shorted to another electrode. For Monkey E, each channel was treated as a neural unit, and waveforms on the channel were not sorted. For Monkeys N and R, waveforms were sorted online before experiments. This approach yielded 87.5 ± 0.5 units (mean ± standard deviation across sessions) for Monkey E, Array 1; 109.6 ± 0.9 units for Monkey E, Array 2; 64.3 ± 13.5 units for Monkey N; and 120.75 ± 48.9 units for Monkey R.

### Kinematic recordings

During reaching tasks, the 3-D position of the hand contralateral to the recording array was tracked using an infrared LED marker attached to the hand (Monkey E, 120Hz sampling rate, PhaseSpace; Monkeys N and R, 60Hz sampling rate, Optotrak 3020, Northern Digital Instruments). Hand position was also recorded for some BCI and isometric force sessions for Monkey E.

During Monkey E’s isometric force task, forces exerted by the hand contralateral to the recording array were recorded using a force transducer (Mini 40, ATI Industrial Automation). The animal held a cylindrical metal handle which was fixed to the force transducer. Hand forces were also recorded for some of Monkey E’s BCI sessions and all tasks of Monkey E’s multiple-tasks paradigm. All force thresholds for all tasks are expressed relative to the amount of force exerted on the force transducer by the hand when the arm was at rest.

### Tasks

In this study we conducted four types of tasks: BCI, isometric force, reaching, and a ‘multiple tasks’ paradigm, and each is detailed below. For all experiments, the monkey sat in a primate chair in a darkened room facing a mirror ∼8cm in front of the eyes which reflected a computer monitor displaying task events. Monkey E was head-fixed, but Monkeys N and R were not.

During reaching tasks, the working arm was not visible to the animal, as it moved in the space behind and below the mirror.

Postural manipulations for all tasks other than the reaching tasks (i.e., the ‘delayed center-out reaching’ task and the ‘center-out reaching’ task) are described relative to a ‘neutral’ posture, in which the upper arm rested on the side of the body, and the forearm was perpendicular to the upper arm, aligned with the sagittal axis (see **Figure 2a**, central posture, for illustration).

### BCI Task

#### Task flow

Monkeys performed an eight-target center-out BCI task by volitionally modulating recorded neural activity to control the position (Monkey E) or velocity (Monkeys N and R) of a computer cursor on a screen. Each experimental session consisted of several blocks. In each block, the arm contralateral to the recording array was fixed in a new posture, and a new decoder was calibrated (see procedure below). A separate decoder was calibrated for each posture so that the animal did not need to produce neural activity appropriate for a different posture’s decoder for task success (**Figure S3**). The monkey used the new decoder to complete between 100 and 200 trials of the center-out task. The arm ipsilateral to the recording array was lightly restrained throughout.

For Monkey E, fine postural manipulations were used. To accomplish this, Monkey E’s contralateral arm was restrained in one of two mechanical devices. Each device allowed for arm posture to be changed by rotating the arm about one joint. The device could then be locked, restraining the arm in the new posture. The first device was used to rotate the forearm about the shoulder in the transverse plane in 15-degree increments. The second was used to rotate the forearm about the elbow in the sagittal plane in 15-degree increments. When using either device, the monkey grasped a handle that was attached to a force transducer so that forces exerted at the hand could be measured during BCI control. Measured forces, motion tracking of the hand, and observed live video feed from the experiment showed that exerted forces and hand movements during BCI control were minimal.

For Monkeys N and R, coarser postural manipulations were used. Arm posture was changed by restraining the arm contralateral to the recording array in a brace placed in different configurations (45-degree rotation of the forearm about the shoulder in the transverse plane and 90-degree abduction of the upper arm about the shoulder in the frontoparallel plane). Hand and arm movements were observed to be minimal during the experiment, but we did not measure them for Monkeys N or R.

### BCI calibration

We used two different types of BCI decoders across animals in this study. For each animal, we used the type of decoder that the animal was familiar with from other experiments prior to this study. The fact that we found similar results regardless of the type of decoder used further strengthens our results.

We begin by describing the calibration procedure for Monkey E, who used a position-based decoder. At the beginning of each session, we estimated the root mean square voltage of the signal on each electrode while the monkey sat calmly in a darkened room. We then began a calibration block of trials, and we repeated the calibration procedure after each postural change.

We used a closed-loop calibration procedure similar to Sadtler et al., 2014^50^. The procedure consisted of five 32-trial blocks of center-out movements (four trials to each of the eight targets, chosen pseudo-randomly). After each block, a new decoder was calibrated that gave the animal increased control of the BCI. The final decoder produced from this procedure was then used by the animal to perform experimental tasks. We describe the calibration procedure in greater detail below.

The first block was an observation block during which we recorded neural activity while the animal observed a computer cursor (circle, 4mm radius) moving from the center target (circle, 15mm radius) to a peripheral target (circle, 10mm radius, 90mm away from center) at a constant velocity (0.15 m/s). Each trial began with a 250ms period during which the screen was blank, and the animal’s hand rested on the handle attached to the force transducer. Next, there was a 300ms ‘freeze period’ during which the cursor, center target and peripheral target were displayed, and the cursor was frozen at the center target location. The cursor then began moving to the peripheral target. After it reached the peripheral target, the animal received a water reward. Data from these observation trials were used to calibrate an initial decoder.

For blocks 2 to 5, the animal performed a BCI center-out task by using the decoder calibrated on trials from the previous block(s) to control the position of the cursor. Each trial began with a 250ms period during which the screen was blank, and the animal’s hand rested on the handle attached to the force transducer. Next, there was a freeze period (500ms) during which the central target and peripheral target appeared, but the cursor was not displayed. After the freeze period, the central target disappeared, and the cursor appeared at a location decoded from neural activity during the freeze period. The animal then had 4s to acquire the peripheral target. If the target was successfully acquired within the time limit, a water reward was given.

In the 2^nd^ block, the initial decoder calibrated on observation trials was used. We restricted movement of the cursor so that it moved in a straight line towards the target (that is, any cursor movement perpendicular to the target was scaled by a factor of 0). For the 3^rd^ block, the decoder was calibrated on the trials from the 2^nd^ block only, and perpendicular movements were scaled by 0.25. For the 4^th^ block, the decoder was calibrated on trials from blocks 2 to 3, and perpendicular movements were scaled by 0.5. For the 5^th^ block, the decoder was calibrated on trials from blocks 2 to 4, and perpendicular movements were no longer restricted (i.e., scaled by 1). The final decoder resulting from the calibration procedure was calibrated on trials from blocks 2-5.

Throughout all calibration blocks, forces at the hand were monitored, and excessive forces triggered a trial failure. Excessive forces were defined as upward or downward forces greater than 3N.

Threshold crossings from calibration trials were binned in non-overlapping 45ms bins. Neural activity from the beginning of the freeze period until target acquisition for each trial was used to fit a Gaussian Process Factor Analysis (GPFA) model to estimate low-dimensional latent states from the high-dimensional neural recordings^51^. These latent state estimates were then used to determine a linear mapping from low-dimensional latent states to cursor position in a procedure described below.

GPFA is a latent variable model that extracts smooth, low-dimensional neural trajectories from simultaneous recordings of many neural units. A neural trajectory summarizes the time evolution of the activity of the recorded neural units. Like factor analysis (FA), GPFA describes the high-dimensional neural activity using a lower-dimensional set of ‘factors,’ which represent the inferred latent state. However, unlike in FA, in GPFA, these latent states are related through Gaussian processes, which embody the notion that trajectories should be smooth. This unifies the typical steps used in extracting neural trajectories (dimensionality reduction and smoothing) into a common probabilistic framework.

GPFA defines a linear-gaussian relationship between a high-dimensional vector of spike counts, *u*_:,*t*_ ∈ ℝ^*q*×1^, (*q* simultaneously recorded channels at timestep *t* = 0,1, …, *T*), and the corresponding low-dimensional latent state at time *t*, *Z*_:,*t*_ ∈ ℝ^*p*×1^ (*p* latent dimensions, *p* < *q*):

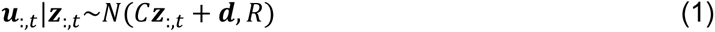

where *C* ∈ ℝ^*q*×*p*^, *d* ∈ ℝ^*q*×1^, and *R* ∈ ℝ^*q*×*q*^ are model parameters to be estimated. As in factor analysis, the covariance matrix *R* is constrained to be diagonal, where the diagonal elements are the independent noise variances of each neural unit.

The latent states at different time points *Z*_:,*t*_ are related through Gaussian processes (GP’s). We define a separate GP for each dimension of the latent state indexed by *i* = 1,2, …, *p*

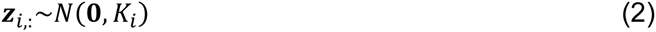

where *K*_*i*_ ∈ ℝ^*T*×*T*^is the covariance matrix for the *i*^th^ GP. Here we chose the squared exponential covariance function as in Yu et al., 2009^51^. Model parameters are fit by the EM algorithm to maximize the probability of the recorded neural activity as described in Yu et al., 2009^51^.

To enable online BCI control, we used a causal implementation of GPFA^52^ in which the estimated latent state at time *t*, *Z*^_:,*t*_, depends only on the current and previous 6 time bins of neural activity (i.e., 7 total bins). We first defined

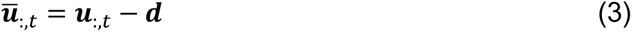

We then concatenated them into a vector *u̅* ∈ ℝ^(7∗*q*)×1^:

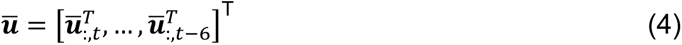

Finally, we used a smoothing kernel, *M* ∈ ℝ^*p*×(7∗*q*)^, to compute our estimate of the latent state at time *t*, *Ẑ*_:,*t*_:

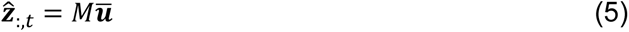

The smoothing kernel, *M*, describes the influence of past spiking activity on the latent state at time *t*, and is determined using *C*, *R*, and *K*_*i*_ for *i* = 1, …, *p*. We used *p* = 10 dimensions for all experiments, as this has been found to capture the majority of shared variability in M1 activity during BCI control^50^.

We then determined a linear mapping from latent state estimates, *Ẑ*_:,*t*_, to 2-D cursor position, *p*_:,*t*_ ∈ ℝ^2×1^:

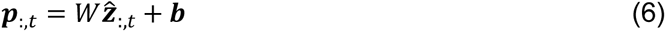

where *W* ∈ ℝ^2×*p*^ and *b* ∈ ℝ^2×1^ are the parameters of the mapping. To estimate *W* and *b*, we used a subset of latent state estimates from each trial and their corresponding intended cursor positions. Specifically, we used latent state estimates from the first 135ms of the freeze period (three bins) and last 225ms before target acquisition (five bins) from each trial (i.e., eight total bins per trial). The intended cursor positions corresponding to these latent state estimates were as follows: for the three bins at the beginning of the freeze period, the intended cursor position was taken to be the screen center in mm, (0,0)^T^. For the 5 observations just before target acquisition, the intended cursor position was taken to be the target location in mm (e.g., for trials to the rightward target, (0,90)^T^).

For Monkeys N and R, each decoder was calibrated by mapping neural activity collected during calibration trials to cursor velocity using the population vector algorithm (PVA, Monkey N)^53,54^ or the optimal linear estimator (OLE, Monkey R)^55^. For both, we used a co-adaptive calibration procedure that has been described previously^56^. Briefly, animals completed center-out trials (described below) to each of the eight possible targets while decoder parameters were recursively updated. Linear (cosine) tuning for each neural unit was estimated by regressing the observed firing rates for each trial to target direction. The level of assistance provided to the subject’s cursor control was gradually reduced. After five cycles through the eight targets (∼2min of data collection), the subject was able to reliably control cursor movement, and decoding parameters were fixed for the remainder of the session. The implementation of the PVA and OLE algorithms has been described in detail in previous work from the group^55^. Briefly, we binned threshold crossings (33ms bins, Monkey N; 20ms bins, Monkey R), subtracted baseline firing rates, and smoothed online by averaging the last 5 time bins together. Weighted averages of these firing rates determined the cursor velocity.

### BCI center-out-across task

The BCI task for Monkey E was a 2-D center-out-across task in which the position of the cursor was controlled with neural activity. The animal first acquired a central target, then a peripheral target, then the diametrically opposed peripheral target. Each trial began with a 250ms period during which the screen was blank, and the animal’s hand rested on the handle attached to the force transducer. Next, the central target (circle, 15mm radius) appeared in the center of the screen, and the cursor (circle, 4mm radius) appeared at a location determined by the monkey’s neural activity. The monkey had 4s to acquire the central target with the cursor. After the target was acquired, it disappeared, and the first peripheral target (circle, 15mm radius) appeared in one of eight possible radial locations (90mm from workspace center). The monkey had 4s to acquire the peripheral target. After the first peripheral target was acquired, a second peripheral target immediately appeared (circle, 15mm radius), diametrically opposed to the first peripheral target. The monkey had 4s to acquire that target. If all three targets were acquired within the specified durations, the animal received a water reward. After each trial, there was a blank screen for 300ms before the next trial.

Throughout, forces at the hand were monitored, and excessive forces triggered a trial failure. Excessive forces were defined as upward or downward forces greater than 3N. Failed trials were followed by a 2s timeout before continuing to the next trial. In this work, we only analyzed data from the initial center-out movement (see ‘Neural activity preprocessing’ below).

The posture of the arm contralateral to the recording array was changed in blocks of trials. For some sessions, five arm postures were used (**Figure 2b, d, e**). These postures were the ‘neutral’ posture and four outer postures achieved by rotating the forearm about the shoulder in the transverse plane (30 degrees clockwise, 15 degrees clockwise, 15 degrees counterclockwise, and 30 degrees counterclockwise from the neutral posture). For other sessions, four arm postures were used (**Figure 2f**). Two of these were achieved by rotating the forearm about the shoulder in the transverse plane (30 degrees clockwise and 30 degrees counterclockwise from the neutral posture). The other two postures were achieved by rotating the forearm about the elbow (30 degrees clockwise and 30 degrees counterclockwise from the neutral posture).

### BCI center-out task

The BCI task for Monkeys N and R was a 2-D center-out task in which the velocity of the cursor was controlled with neural activity. The animal moved the cursor (circle, 6mm radius) from the center of the screen to a peripheral target on each trial. At the beginning of each trial, a central target appeared (circle, 6mm radius) and the cursor position was reset to the center of the screen. The animal was required to hold the cursor within the central target for 50ms (Monkey N) or 200ms (Monkey R). Next, the central target disappeared and a peripheral target (circle, 6mm radius, Monkey N; 5mm radius, Monkey R) appeared 85mm away. The monkey had 3s to acquire the peripheral target and was required to hold it for 200ms (Monkey N) or 100ms (Monkey R) to receive a water reward. After each trial, there was a blank screen for 500ms before the next trial.

The posture of the arm contralateral to the recording array was changed in blocks of trials. For Monkey N, three postures were used in each session: (1) the neutral posture, (2) a 45-degree rotation of the forearm about the shoulder in the transverse plane and (3) a 90-degree abduction of the upper arm about the shoulder in the frontoparallel plane (**Figure 2g**). For Monkey R, two postures were used in each session. In one session, (1) the neutral posture and (2) a 45-degree rotation of the forearm about the shoulder in the transverse plane were used. In another session, (1) the neutral posture and (2) a 90-degree abduction of the upper arm about the shoulder in the frontoparallel plane were used (**Figure 2h**).

### Isometric force task

Monkey E completed an isometric force task in which an upward or downward force was exerted on the handle attached to the force transducer. Forces were exerted by the hand contralateral to the recording array. Forces were mapped to the position of a computer cursor that was constrained to move on a vertical line, so that only upward or downward components of exerted forces contributed to the cursor motion. The position of the hand used to exert forces was recorded using the motion tracking system. The ipsilateral hand was lightly restrained.

At the beginning of each trial, the cursor (circle, 4mm radius) appeared in a location dictated by the current level of force exerted on the handle, and a central target (square, 10mm side length) appeared, corresponding to the amount of force produced by the hand lightly resting on the bar. The animal had to acquire the central target within 4s and hold it for 350ms. Once the central target was acquired, it disappeared, and a peripheral target (square, 10mm side length) appeared 38mm above or below the location of the central target. The animal then had to acquire the peripheral target within 4s and hold it for 350ms to receive a water reward. To acquire the upward or downward peripheral target, the animal had to exert 4.75±0.62N of upward or downward force. There was a 300ms period before the start of the next trial.

The posture of the arm used to exert forces was changed in blocks of trials. Five arm postures were used (**Figure 4a**). These postures were the ‘neutral’ posture and four outer postures achieved by rotating the forearm about the shoulder in the transverse plane (30 degrees clockwise, 15 degrees clockwise, 15 degrees counterclockwise, and 30 degrees counterclockwise from the neutral posture).

### Reaching Task

#### Center-out reaching task

The center-out reaching task for Monkeys N and R was a 2-D, eight-target task. Animals made unconstrained reaches in 3-D space, but only the vertical and horizontal components of those reaches were mapped to cursor motion. Hand position was recorded as described above and mapped to the location of the computer cursor (circle, 5mm radius). The mapping was calibrated such that 1cm of hand displacement in the frontoparallel plane corresponded to 1cm of cursor movement. At the beginning of each trial, a central target appeared (circle, 5 mm radius), and the animal had 3s to acquire it with the cursor. Once it was acquired, the animal had to hold it for a variable duration ‘center hold period’ (0-100ms, uniformly distributed). Next, go cue was indicated by the disappearance of the central target and appearance of a peripheral target (circle, 5mm radius) located 60mm away from the central target. The animal had 1000ms to acquire the peripheral target. Finally, the peripheral target was held for a variable length period (200-400ms, uniformly distributed). If the peripheral target was acquired within 1000ms and held for the entire target hold period, the animal received a water reward.

The initial posture of the arm was changed in blocks of trials by changing the initial location of the hand in the frontoparallel plane. Reaches were matched in direction and distance from each initial hand position. Visual feedback was matched across postures so that the instructed initial hand position always corresponded to the center of the screen. There were 2 initial hand locations. These were located (-51.2, 34.7) and (51.2, -34.7) mm (x, y) away from the center of the animal’s workspace (i.e., the initial hand locations were separated diagonally by 123.7mm). The ipsilateral arm was lightly restrained during the task.

### Delayed center-out reaching task

The reaching task for Monkey E was a 2-D, eight target delayed center-out task. Hand position was recorded and mapped to the location of the computer cursor as described above. At the beginning of each trial, the central target appeared (circle, 12mm radius), and the animal had 5s to acquire it with the cursor. Once it was acquired, the animal had to hold it for a variable duration (450ms, 40% of trials; 550ms, 40%; 1000ms, 20%). If the cursor left the central target during the center hold period, the animal could bring the cursor back into the central target and restart the hold period. Once the center hold period was complete, the peripheral reach target appeared (circle, 12mm radius, 80mm from center target) and a variable-length delay period began (25ms, 500ms, 750ms, or 1000ms, equal probability) during which the animal continued to hold the cursor within the center target. Next, the go cue was indicated by the disappearance of the central target, after which the animal had 1500ms to acquire the peripheral target. Finally, the animal was required to hold the cursor in the peripheral target for a variable duration (500ms, 66% of trials, 1000ms, 34%). After a successful trial, there was a 300ms pause before the next trial. If the trial was failed, there was a 650ms penalty. 2.5% of trials were ‘catch trials’, which ended with a reward after the delay period if the animal held the cursor in the central target for the duration of the delay period. These trials were not analyzed but helped to keep the animal motivated to not leave the central target before the go cue was given. The delay period was not necessary for this study, which focuses on peri-movement activity. It was included to study the effect of posture on neural activity during the delay period, which will be explored in a future study.

The initial posture of the arm was changed in blocks of trials by changing the initial location of the hand in the frontoparallel plane. Reaches were matched in direction and distance from each initial hand position. Visual feedback was matched across postures so that the instructed initial hand position always corresponded to the center of the screen. There were 7 initial hand locations: a central location, a row of four locations above the central location, and a row of two locations below the central location. The coordinates of each location relative to the central location in mm (x, y) were as follows: (-120,40), (-40,40), (40,40), (120,40), (0,0), (-40,-40), and (40,-40). The ipsilateral arm was lightly restrained during the task.

### Multiple tasks paradigm

Monkey E also completed a ‘multiple tasks’ paradigm in which multi-posture reaching, BCI, and isometric force tasks were all completed within a single session. This paradigm was designed to compare the influence of posture on neural activity across tasks. Therefore, the initial postures of the arm were matched across tasks. To achieve this, each trial of the paradigm began with the animal holding the handle fixed to the force transducer with the hand contralateral to the recording array. This hand was used to exert forces in the isometric force task and to make reaches in the reaching task. The posture of the arm holding the handle was changed in blocks of trials. The same three initial arm postures were used for all tasks (**Figure 5a**). These postures were the neutral posture and two outer postures achieved by rotating the forearm about the shoulder in the transverse plane 30 degrees clockwise or counterclockwise from the neutral posture. The isometric force task was carried out exactly as described in the ‘isometric force task’ section above.

The BCI task was an 8-target center-out-across task similar to that described above with a few minor differences. Each trial began with a 250ms period during which the screen was blank, and the animal’s hand rested on the handle attached to the force transducer. Next, a central target (circle, 15mm radius) and peripheral target (circle, 15mm radius, 90mm away from center) appeared and a 500ms ‘freeze period’ began during which the cursor was not displayed.

However, neural activity recorded during the freeze period was used to determine the position of the cursor in subsequent timesteps. After the freeze period, the central target disappeared and the cursor appeared, and the animal had 4s to acquire the peripheral target. After this, the ‘across’ movement began, and the task proceeded exactly as the ‘BCI center-out-across task’ described above.

The reaching task was a 2-D, three target task. Hand position was recorded and mapped to the location of the computer cursor as described above. At the beginning of each trial, the animal had to hold the handle fixed to the force transducer for 1s. Excessive forces exerted at the hand during this period (upward or downward force > 3.5N) triggered a trial failure. After this, a reach target (circle, 6mm radius) appeared in one of 3 possible locations, and the animal had 3s to acquire it with the cursor. The target locations for each posture were (-100,75), (0,125), and (100,75) measured in mm (x,y) from the position of the hand when it held the handle. That is, the directions and distances of the targets from the initial hand position were matched across postures. The target locations were chosen so the animal could make unimpeded reaches from the handle to the target. After acquiring the reach target, the monkey was required to hold it for 300ms. There was no delay period. Visual feedback was matched across postures so that the targets always appeared at the same locations on the screen.

### Experimental sessions

With Monkey E (Array 1), we conducted four sessions of the BCI center-out-across paradigm, three sessions of the isometric force paradigm, and one session of the multiple-tasks paradigm. Only one session of the multiple-tasks paradigm was collected because it was a difficult task for the animal. With Monkey E (Array 2), we conducted one session of the BCI center-out paradigm and three sessions of delayed center-out reaching. With Monkey N, we conducted two sessions of the BCI center-out paradigm and five sessions of the center-out reaching paradigm. With Monkey R, we conducted two sessions of the BCI center-out paradigm and two sessions of the center-out reaching paradigm.

### Data analysis

#### Kinematic data preprocessing and analysis

Here we describe how we determined the movement onset times in the reaching tasks. These times were used to select the analysis window for each trial of the reaching tasks as described in the ‘Neural activity preprocessing’ section below.

Hand position signals were first processed as in Smoulder et al., 2023^45^. In brief, position measurements were smoothed with a zero-phase low pass Butterworth filter (total 8^th^ order, cutoff frequency 15Hz). Velocities were computed by taking the first difference of the position signals, dividing by the time difference between samples, and assigning each velocity sample a timestamp that was the midpoint of the timestamps for the two position samples used. We then used spline interpolation to upsample position and velocity measurements to 1000Hz.

To determine the movement onset time for each trial in the reaching tasks, hand speed was calculated at each time step using the horizontal and vertical components of hand velocity. We then determined the peak speed for the trial as the maximum hand speed between go cue and target acquisition. Finally, movement onset was taken as the earliest time after go cue at which hand speed reached 20% of the peak speed for the trial.

Only successful trials from each task were analyzed. To ensure we were analyzing trials with consistent behavior, we removed any trials with abnormally long movement epochs from each session (95^th^ percentile and above for each session; movement epoch is the time from go cue to target acquisition).

### Neural activity preprocessing

For all tasks, we focused on the early execution epoch, defined for each task below, when posture and goal information were simultaneously present in neural activity. For BCI tasks, we analyzed the 200ms period beginning 50ms after go cue/target onset. We began 50ms after target onset, as visual information takes more than 50ms to reach motor cortex^57^. We ended 250ms after target onset to exclude any corrective cursor movements that occurred later in the trial. For isometric force tasks, we also analyzed the 200ms period beginning 50ms after go cue/target onset. For reaching tasks, we analyzed the 200ms period before movement onset. The ends of the analysis windows for the isometric force and reaching tasks were chosen to exclude any time-varying proprioceptive signals produced by muscle contraction.

Neural spike times were convolved with a Gaussian kernel (25ms standard deviation) to produce estimates of firing rate versus time. These firing rates were sampled every 25ms during the analysis window for each task. The sampled firing rates for each electrode were z-scored separately to prevent electrodes with high firing rates from overly influencing population-level results. The mean and standard deviation used for z-scoring each electrode were computed using all sampled firing rate estimates taken when animals were engaged in the task (center hold period to the end of trial) for all trials that met the inclusion criteria (outlined in the ‘Kinematic data preprocessing and analysis’ section above). Electrodes for which this mean firing rate was < 3 Hz were excluded from all neural analyses.

### Principal component analysis (PCA)

To produce the PCA visualizations in **Figure 2d** and **Figure S2**, we first formed condition-averaged neural trajectories for each posture and target combination. We then concatenated these condition-averaged trajectories into an *N* × *CT* matrix, where *N* is the number of recorded units, *C* is the number of conditions (i.e., the number of postures times the number of targets), and *T* is the number of timesteps in each trajectory. PCA was performed on this matrix to identify the directions of greatest variance within the *N*-dimensional population activity space.

### Targeted dimensionality reduction

To create the ‘targeted dimensionality reduction’ (TDR) visualizations in **Figure 2** (**e**-**h**) and **Figure 4** (**b**&**e**), we used demixed principal components analysis (dPCA)^41^ [https://github.com/machenslab/dPCA]. The default dPCA parameters were used, including the optimization procedure to find the regularization factor.

dPCA is a linear dimensionality reduction technique which identifies neural dimensions that explain variance related to individual experimental variables. To do this, dPCA first decomposes the activity from each neural unit into marginalizations in an ANOVA-like manner. It then identifies neural dimensions which explain variance in groups of these marginalizations. We explain below how to compute the marginalization groups used to identify posture and goal dimensions in Figure 2 (e-h) and Figure 4 (b&e).

Consider the condition-averaged activity recorded from one neural unit for one experimental session. We first subtracted the overall mean activity for the unit (across all trials and timesteps) from its condition-averaged activity at each timestep. We next collected the activity into a 3- dimensional tensor, *x*_*t,gp*_ ∈ ℝ^*T×G×P*^, where *T* is the number of timesteps, *G* is the number of goals (i.e., targets), and *P* is the number of postures. For this work, the relevant marginalizations of this tensor were the *T* × 1 ‘time marginalization,’ formed by averaging over all variables except for time; the *T* × *P* ‘posture marginalization,’ formed by averaging over goal, then subtracting the time marginalization from each column (to remove time effects); and the *T* × *G* ‘goal marginalization’, formed by averaging over posture, then subtracting the time marginalization from each column (to remove time effects).

After computing these marginalization groups, we combined them across units to form the input for dPCA. dPCA was used to identify time, posture, and goal dimensions that sought to explain variability from their respective marginalization groups while minimizing variance captured from other marginalizations. For visualization, we show the top two goal dimensions and the top posture dimension in a 3-D space after orthonormalizing them using QR decomposition.

### Cross-projection alignment test

We were interested in whether the subspaces modulated by posture- and goal-related neural signals were misaligned (**Figure 3**, **Figure 4**, and **Figure S5**). We therefore first identified posture- and goal-related components of neural activity (see procedure below). We then identified the subspaces modulated by each, which we refer to as the posture and goal subspaces, respectively. To quantify the alignment of these subspaces, we employed the cross-projection alignment test in Elsayed et al., 2016^19^. This test measures the alignment of two subspaces by projecting the same neural activity into each subspace, then comparing the amount of variance captured by each subspace. If the subspaces are aligned, a similar amount of variance will be captured by each subspace. However, if the subspaces are not aligned, they will capture different amounts of variance. For our work, this test is preferrable to measuring the angles between subspaces, because it takes into account the variance of the neural activity along different dimensions. To measure the alignment of the posture and goal subspaces, we projected the posture component into the posture and goal subspaces and compared the amount of variance captured by each. For completeness, we repeated this procedure with the goal component.

We first identified the 10-D space that captured the greatest variance in the trial-averaged neural activity for each session using PCA (see ‘Principal component analysis’ above). This 10- D space accounted for the majority of the variance in the trial-averaged neural activity (90 ± 4%, mean ± s.d. across datasets). Neural activity from each trial was projected into this 10-D space before further analysis. This step ensures (1) that our analyses focus on aspects of the activity with greatest variance, and (2) that when comparing the variance captured by a subspace to that captured by random subspaces (procedure below), the random subspaces are drawn from the space occupied by neural activity. We then divided the trials from each task condition (e.g., posture 1, target 1) into two equally-sized folds for cross-validation. We included only postures for which there were at least 10 trials to each target direction (90% of postures across datasets). This ensured that there were at least 5 trials for each condition in each of the 2 cross-validation folds.

We next measured the amount of posture-related variance captured by the goal subspace. Using trials from the first fold, we identified the posture component by computing the *T* × *P* posture marginalization for each of the *N* neural units as described in the ‘Targeted dimensionality reduction section’ above. The columns of each of the *N* posture marginalizations were concatenated vertically, forming *N* column vectors of size *TP* × 1. These column vectors were then concatenated horizontally and transposed, forming the *N* × *TP* posture component. To identify the goal subspace, we first used an analogous procedure to compute the *N* × *TG* goal component on trials from the second fold. We then applied PCA to the goal component to identify the goal subspace. We used PCA instead of dPCA to identify the subspace, since the loss function of dPCA can encourage the identification of orthogonal subspaces for each task variable^41^. We kept only the top 2 PCs of the goal subspace, as these accounted for the majority (85 ± 7%, mean ± s.d. across datasets) of the variance in the goal component. We projected the posture component from the first fold into the goal subspace from the second fold and measured the amount of variance captured. We then repeated this entire procedure using the second fold to identify the posture component and the first fold to identify the goal subspace. The amount of posture-related variance captured by the goal subspace was taken to be the mean across the two folds.

We used the same technique to measure the variance of each component in each subspace. Specifically, we measured the amount of posture-related variance captured by the posture subspace, goal-related variance captured by the posture subspace, and goal-related variance captured by the goal subspace. When identifying the posture subspace, we again kept only the top 2 PCs, as they accounted for the majority (92 ± 8%, mean ± s.d. across datasets) of the variance in the posture component.

To compute confidence intervals for these measurements, we again used neural activity from the second fold to identify posture and goal subspaces. We then used bootstrap resampling to produce 10,000 resampled versions of the first fold. For each of these bootstrap resamples, we computed posture and goal components, projected each into the posture and goal subspaces, and measured the amount of variance captured. We repeated this procedure using the first fold to identify subspaces and the second fold for projection. This produced 20,000 bootstrapped estimates of variance captured in each component by each subspace. These bootstrapped estimates were then combined across sessions for each monkey and task. For each monkey and task, the 95% confidence interval was taken as the 2.5th percentile to the 97.5th percentile of this combined bootstrapped distribution. To assess whether variance captured in each component by the posture and goal subspaces were significantly different (p<0.05), a paired, two-tailed bootstrap test was performed using the session-combined bootstrap distributions. For example, the variance in the posture component captured by the posture and goal subspaces on each bootstrap resample were paired.

To estimate the distribution of variance in each component captured by random subspaces, we drew 10,000, 2-dimensional random subspaces in the 10-D PC space. Drawing these spaces from within the 10-D space explaining the most variance in the activity (identified with PCA) ensured that random subspaces are drawn from dimensions where neural activity resides. Each random subspace was constructed by drawing two vectors in the 10-D PC space and orthonormalizing them. Each component of each vector was drawn from a standard normal distribution. We projected the posture and goal components from each of the two folds into these 10,000 subspaces and measured the amount of variance captured. This procedure produced 20,000 variance captured measurements for each component. The measurements for each component were then combined across sessions and monkeys for each task. This resulted in one distribution for each component for each task.

### Assessing the consistency of the posture subspace across tasks

To assess the consistency of the posture subspace across tasks (**Figure 5**), we tested whether it was possible to identify a single subspace from which arm posture could be read out regardless of task. To test this, we analyzed neural activity from the multiple tasks paradigm. In this paradigm, Monkey E performed three tasks (BCI, isometric force, and reaching) from three initial arm postures, which were matched across tasks. We first visualized neural activity from each task to determine whether it was clustered by posture in a posture subspace identified across all tasks. If so, this would suggest a consistent posture subspace across tasks. We then asked whether arm posture could be accurately classified from neural activity using a single classifier across tasks. This would further indicate a consistent posture subspace across tasks.

For all analyses in Figure 5, we averaged the z-scored firing rates over all timesteps in the analysis window to produce a single *N* × 1 firing rate estimate per trial, where *N* is the number of recorded neural units. The analysis windows used for each task of the multiple tasks paradigm were the same as those used when analyzing individual task paradigms. Neural activity from all trials was then collected into a matrix of size *N* × *K*, where *K* was the number of trials in the dataset. PCA was performed on this matrix to identify the top principal components within the *N*-dimensional population activity space. Each trial of neural activity was then projected onto the top 10 principal components, producing a matrix of size 10 × *K*. Reducing the dimensionality of the neural activity in this way allowed us to better estimate the covariance matrix of the activity with limited trials.

To create the visualizations in **Figure 5b**, we used linear discriminant analysis (LDA) to identify neural dimensions that separated the 10-D neural population activity by posture, regardless of task or target direction. This was done only to visualize the data, and cross-validated classification analyses were performed subsequently (see below). The class-specific means for LDA were computed by taking the mean of the activity from each posture. The covariance matrix for LDA was computed by first computing the covariance matrix for each posture separately, then averaging them across postures. We subsampled trials so that there were (1) equal numbers of trials from each task in each posture and (2) approximately equal numbers of trials from each target within each task-posture combination.

We next used maximum likelihood estimation to classify neural activity from each trial by posture (**Figure 5c**). We used the statistical model from LDA, in which each posture has its own mean, and all postures share a single covariance matrix. All classification analyses were cross-validated. We performed three separate classification analyses: ‘within task,’ ‘train on one task, test on others,’ and ‘train on all tasks.’ The ‘within task’ analysis assessed how well we could classify posture within each task, and the other analyses assessed how well classifiers generalized across tasks. Our metric for measuring classifier performance was the percentage of correct classifications.

For the ‘within task’ analysis, we trained a classifier using trials from one task and tested it using held-out trials from the same task. This was done separately for each of the three tasks. We used 5-fold cross-validation. We subsampled trials within each task so that there were equal numbers of trials from each target direction in each posture. We then subsampled from the remaining trials such that the number of trials in the training set for each task was equal.

For the ‘train on one task, test on others’ analysis, we tested each classifier from the ‘within task’ analysis on the two tasks it was not trained on. In this setting, the training and testing trials correspond to different tasks, so there was no need for cross-validation. However, to fairly compare to the ‘within task’ setting, we used the same training and test partitions. After training each classifier on four folds of data, we tested it on the corresponding test fold of another task (e.g., trained on folds 1-4 of isometric force trials and tested on 5^th^ fold of BCI trials).

For the ‘train on all tasks’ analysis, we trained a single classifier on trials from all tasks then tested it on each task separately. To fairly compare to the ‘within task’ setting, we again used the same training and test partitions. We formed training sets using trials from 4 folds of each task, then tested on the remaining fold from each task (e.g., we trained on trials from folds 1-4 of BCI, isometric force, and reaching, and tested separately on fold 5 of BCI, isometric force, and reaching). We subsampled trials so that each training set (1) contained equal numbers of trials from each task and (2) matched the size of the training set used for each individual task in the ‘within task’ analysis.

We performed a final visualization in **Figure 5d** to understand how the posture subspace can be similar across tasks, yet overall neural activity is distinct across tasks. To do this, we grouped trials by task, regardless of posture or target direction. We subsampled trials within each task so that there were equal trial counts for each posture and target combination. We then applied LDA to identify dimensions that separated neural activity by task. Because there were 3 tasks, this procedure resulted in a 2-dimensional subspace. While the neural activity from the 3 tasks was well-separated in the 2-D space returned by LDA, the task means were not evenly-spaced along the first LDA dimension. Therefore, we manually chose one dimension from within this subspace along which data from all three tasks were visibly separated for visualization. The posture dimensions in Figure 5d are the same as those in Figure 5b.

### Assessing the similarity in neural trajectories across postures

To assess the similarity of neural trajectories across postures (**Figure 6, Figure S8**), we asked how well pairs of trajectories from different postures matched if we translated them to bring them into alignment. If a pair matched perfectly after translation, this would indicate that the trajectories had the same shape but started in different locations due to posture-related effects in neural activity. We allowed for translation, but not scaling, because we wanted to focus on the robustness of trajectory shape. Each pair of trajectories corresponded to the same target direction and task, so that any difference in trajectory shape was due only to posture. Our procedure is illustrated in detail for one example comparison in **Figure S8**.

We performed this analysis within the 10-D space identified by applying PCA to the trial-averaged neural activity for each session (see ‘Principal component analysis’ above). Neural activity from individual trials was projected into this 10-D space. We then removed any conditions (i.e., combinations of posture and target, such as posture 1, target 1) containing fewer than 10 trials so that there were adequate trial counts for subsampling.

For each session, we compared every possible pair of trial-averaged trajectories from different postures but the same target direction (e.g., we compared posture 1, target 1, which we refer to as ‘condition 1’, to posture 2, target 1, which we refer to as ‘condition 2’). This led to *P*_*t*_(*P*_*t*_ − 1)/2 comparisons per session, where *P*_*t*_ is the number of postures remaining for the *t*^th^ target after removing conditions with inadequate trial counts, and *T* is the number of targets.

For each comparison, we randomly sampled trials from each condition without replacement to form 2 groups (e.g., condition 1, group 1; condition 1, group 2; condition 2, group 1; condition 2, group 2). Each group contained *N*_*M*_/2 trials, where *N*_*M*_ is the minimum number of trials for any condition in the session. We then computed trial-averaged trajectory for each condition and group (we refer to these trial-averaged trajectories as simply ‘trajectories’ hereafter). Dividing into groups in this way allowed us to estimate a lower bound for the difference between a pair of trajectories (procedure below).

We next assessed how similar trajectories from the two conditions were. To do this, we measured the ‘difference after translation’ between the trajectories from group 1 of each condition. We first brought the trajectories into alignment by translating the trajectory from condition 2, group 1 by the vector connecting its mean (taken across timesteps) to the mean (taken across timesteps) of condition 1, group 1. We then measured the Euclidean distance between these trajectories at each timestep and averaged the distances across timesteps.

We compared the difference after translation to estimates of its lowest possible value and its highest possible value. To estimate the lowest possible value, we measured the difference after translation between trajectories from different groups of the same condition (e.g., condition 1, group 1 and condition 1, group 2). To estimate the highest possible value, we measured the difference *before* translation between the trajectories from group 1 of each condition. This was computed by measuring the Euclidean distance between the trajectories from condition 1, group 1 and condition 2, group 1 at each timestep and averaging the distances across timesteps.

For interpretability of results, we normalized the difference before translation and difference after translation. We did this by dividing each by our estimate of the lowest possible value for the difference after translation. Therefore, after normalization, a value of 1 corresponds to the difference in trajectories arising solely due to analyzing a finite number of trials. We repeated this entire procedure 20 times for each comparison, resampling trials on each repeat. For each metric (i.e., difference before translation and difference after translation), we took the mean across the 20 repeats.

To visualize the results, we plotted the difference before translation versus the difference after translation for each comparison (**Figure 6c**). This format allows for each comparison to be compared to its approximate lowest possible value of 1 (horizontal dashed line) and its highest possible value (diagonal dashed line, corresponding to no decrease in trajectory difference after translation). Due to the finite number of trials analyzed, it was possible for normalized values to be below 1. This occurred when the trial-averaged trajectories from across conditions were more similar in shape than those from different groups within the same condition.

We also computed the mean percentage reduction in trajectory difference (before versus after translation) across all comparisons for each animal. This was reported in the main text accompanying **Figure 6**. When computing this reduction, we de-biased the differences before and after translation for each comparison by subtracting an estimate of the bias from each. To estimate the bias for the difference before translation, we took the mean difference before translation of trajectories from different groups of the same condition. For example, to compute this bias when comparing condition 1 to condition 2, we measured the difference before translation between condition 1 group 1 and condition 1 group 2. We also measured the difference before translation between condition 2 group 1 and condition 2 group 2. We combined all of these measurements across repeats and took the mean. This was our estimate of the bias for the difference before translation for this comparison. An analogous procedure was used to estimate the bias for the difference after translation.

## Statistics for main text figures

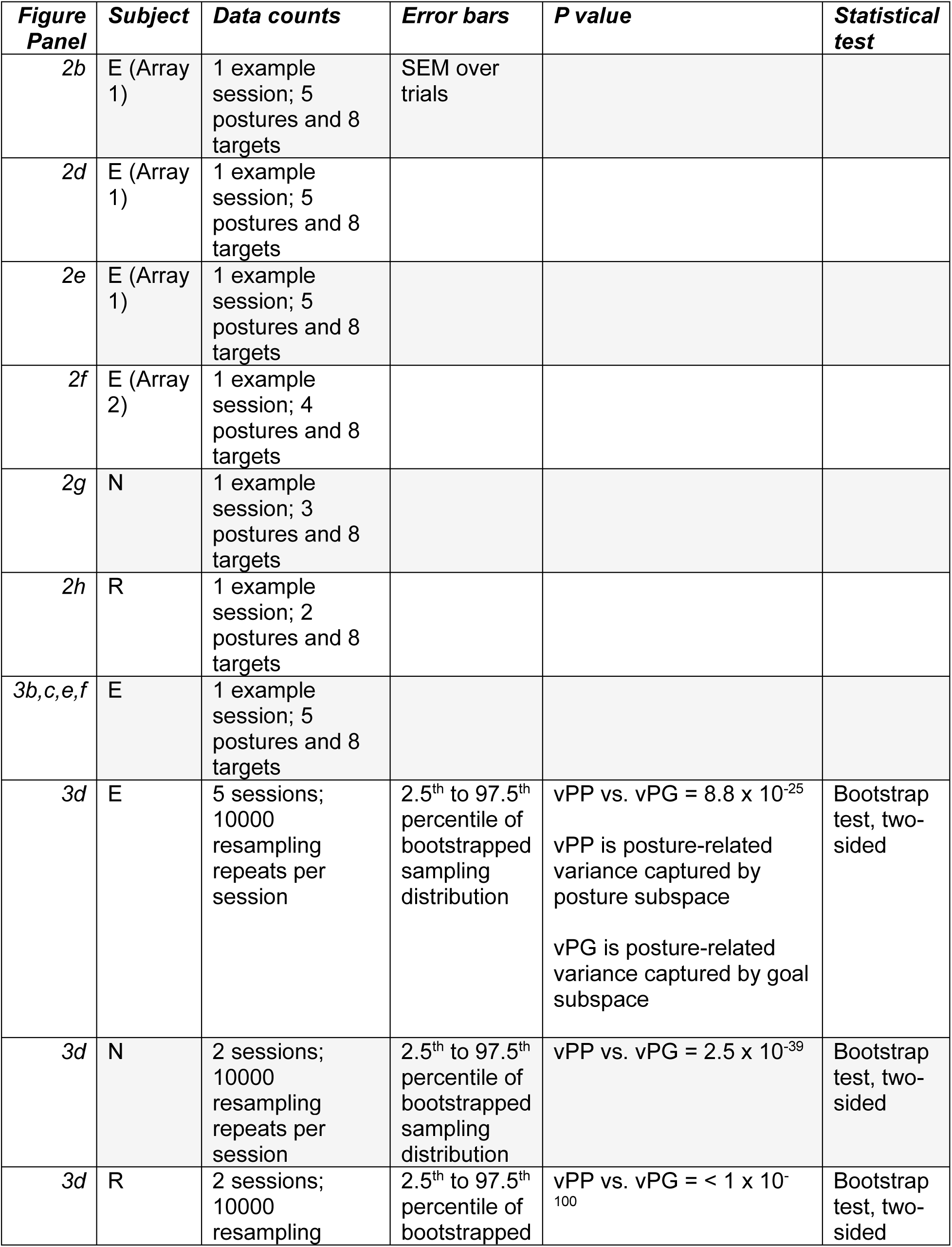
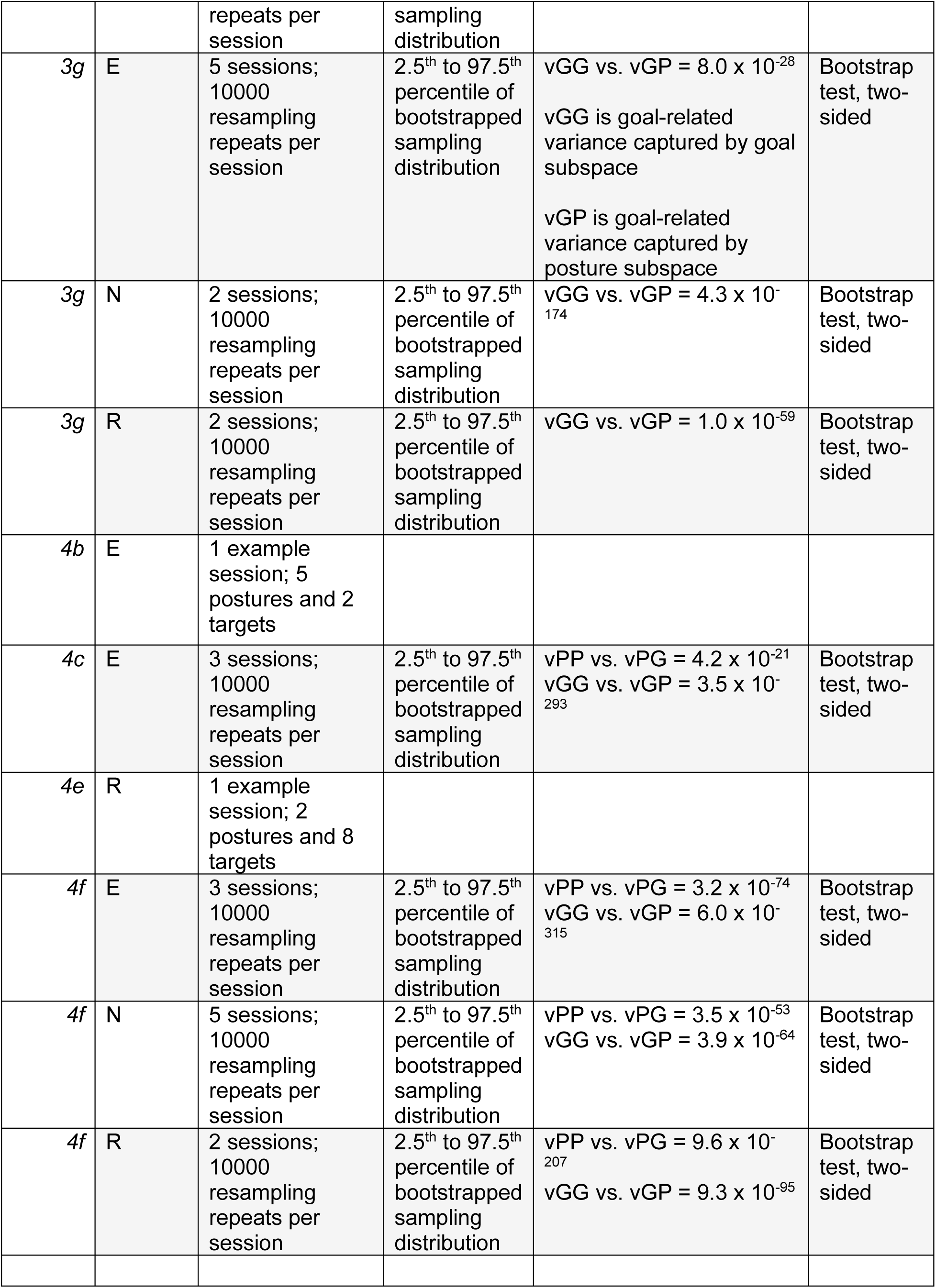

