## Supplementary Material for "A posture subspace in primary motor cortex"

[1]Dept. of Bioengineering, University of Pittsburgh, Pittsburgh, PA 15260, USA
[2]Center for the Neural Basis of Cognition, Pittsburgh, PA 15213, USA
[3]Dept. of Biomedical Engineering, Carnegie Mellon University, Pittsburgh, PA 15213, USA
[4]Dept. of Biomedical and Molecular Sciences, Queen's University, Kingston, Ontario K7L 3N6, Canda
[5]Dept. Electrical and Computer Engineering, Carnegie Mellon University, Pittsburgh, PA 15213, USA
[6]Starfish Neuroscience, Bellevue, WA 98004, USA
[7]Dept. of Physical Medicine and Rehabilitation, University of Pittsburgh, Pittsburgh, PA 15260, USA
[8]Rehab Neural Engineering Labs, University of Pittsburgh, Pittsburgh, PA 15260, USA
[9]Dept. of Neurobiology, Physiology, and Behavior, University of California, Davis, Davis, CA 95616, USA
[10]Neuroscience Institute, Carnegie Mellon University, Pittsburgh, PA 15213, USA
[11]Senior author
[12]These authors contributed equally
[13]Lead contact.

### Supplemental Figures

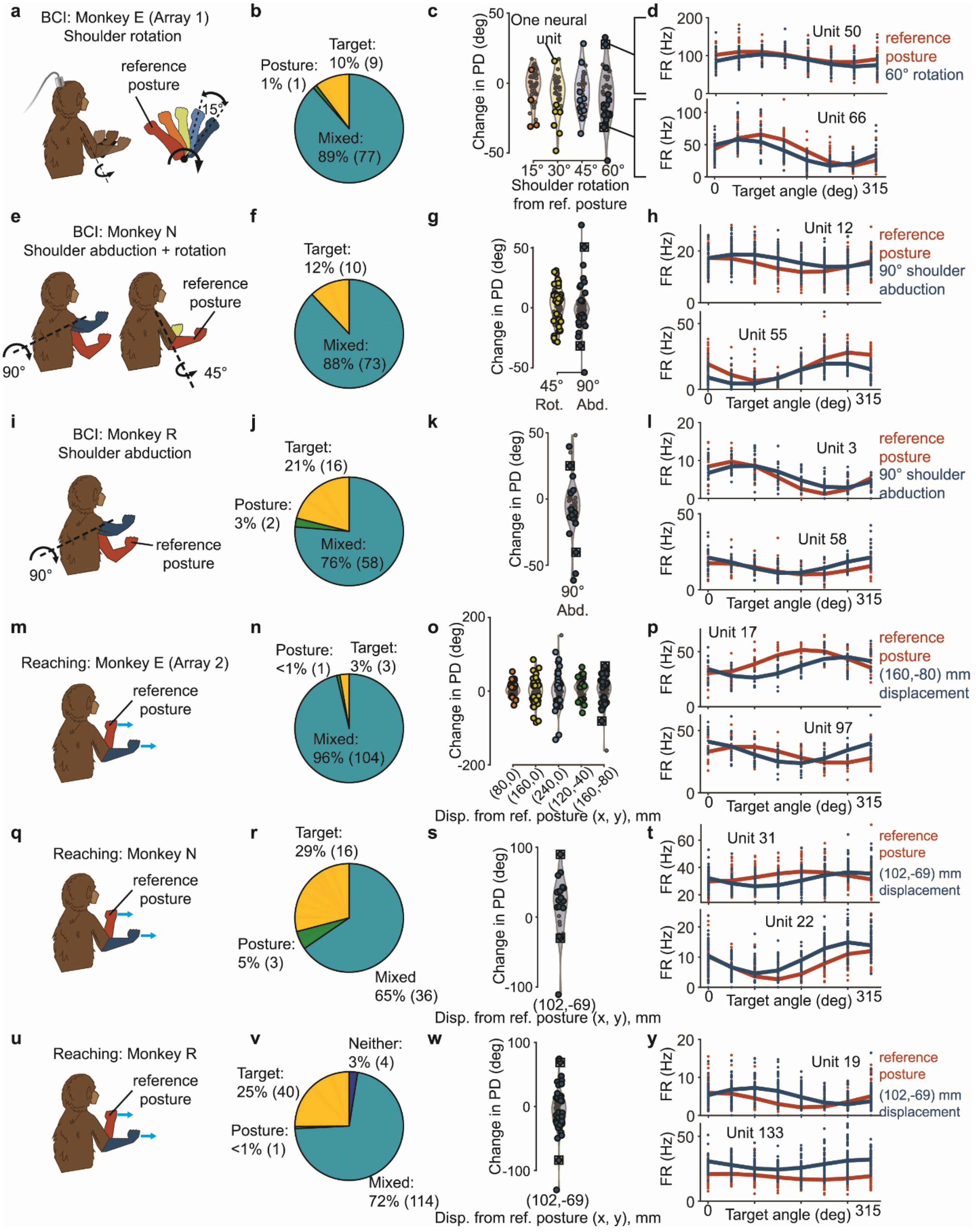

**Supplementary Figure 1| Most individual units were tuned to target and posture, and many exhibited changes in preferred direction across postures**.

**(a) I**llustration of arm postures used in Monkey E's BCI task (see main text and Methods for detailed description). Signed changes in preferred direction (PD) were measured between the 'reference posture' and each of the other postures, which we refer to as 'comparison postures'. The reference posture was chosen to be at one extreme of the postures used in each session, so that when the comparison posture was at the other extreme, the physical separation (i.e., joint angle or hand displacement) between the reference and comparison postures was maximized. **(b)** Proportion of individual neural units that were significantly tuned to posture, target direction, both (i.e., 'mixed'), or neither for Monkey E's BCI task. For all tasks, we analyzed neural activity from the entire center-out movement (i.e., go cue to peripheral target acquisition), as in previous studies [S1,S2]. Spike counts were taken over the entire analysis window, and no convolution or z-scoring was performed. This resulted in one spike count vector (i.e., an $N \times 1$ vector, where $N$ is the number of recorded neural units) per trial. To determine whether individual units were tuned to target direction and/or posture, we used an unbalanced, 2-way analysis of variance (ANOVA). Units were classified based on whether target, posture, neither, or both had a significant effect (F-test, $p < 0.05$). Most neural units displayed mixed tuning, although some were tuned exclusively to target or posture. **(c)** Distribution of changes in PD of individual units across postures. Larger circles represent units with statistically significant changes in PD across postures, and smaller circles represent units for which changes were not statistically significant. To analyze changes in PD across postures, we first assessed whether units were directionally-tuned in each posture by fitting a cosine tuning model to each unit in each posture [S3]: $f = b_0 + b_1 \sin\theta + b_2 \cos\theta$, where $f$ is the firing rate of the unit, $\theta$ is the reach direction, and $b_0, b_1$, and $b_2$ are the model coefficients. The model was fit to mean firing rates for each reach direction using multiple linear regression. A separate model was fit for each unit in each posture. Units were taken to be directionally tuned in a posture if the regression coefficients of the model were significantly different from zero (F-test, $p < 0.05$). The PD for each unit in each posture was taken to be the value of $\theta$ that maximized the firing rate of the model. When assessing changes in PD across postures, only units that were directionally-tuned in both the reference and comparison posture were analyzed. We used a bootstrapping approach to determine whether measured changes in PD were statistically significant. To do this, we resampled trials for both the reference and comparison postures with replacement, preserving the number of trials for each target in each posture. Next, we computed the PDs for each posture and signed change in PD between the reference and comparison postures using the resampled data. We repeated this procedure 10,000 times to produce a bootstrapped distribution of signed PD changes for the pair of postures. We then assessed whether the mean of the bootstrapped distribution was significantly different from zero ($p < 0.05$). **(d)** Tuning curves for two example units in the reference posture and most extreme comparison posture (60˚ shoulder rotation). The example units chosen for visualization are indicated with squares in (c). Dots indicate firing rate for one trial. Curves show the model fit. Changes in preferred direction between the reference and comparison postures are visible in the example tuning curves. **(e-y)** Same conventions as (a-d) for other animals and tasks.

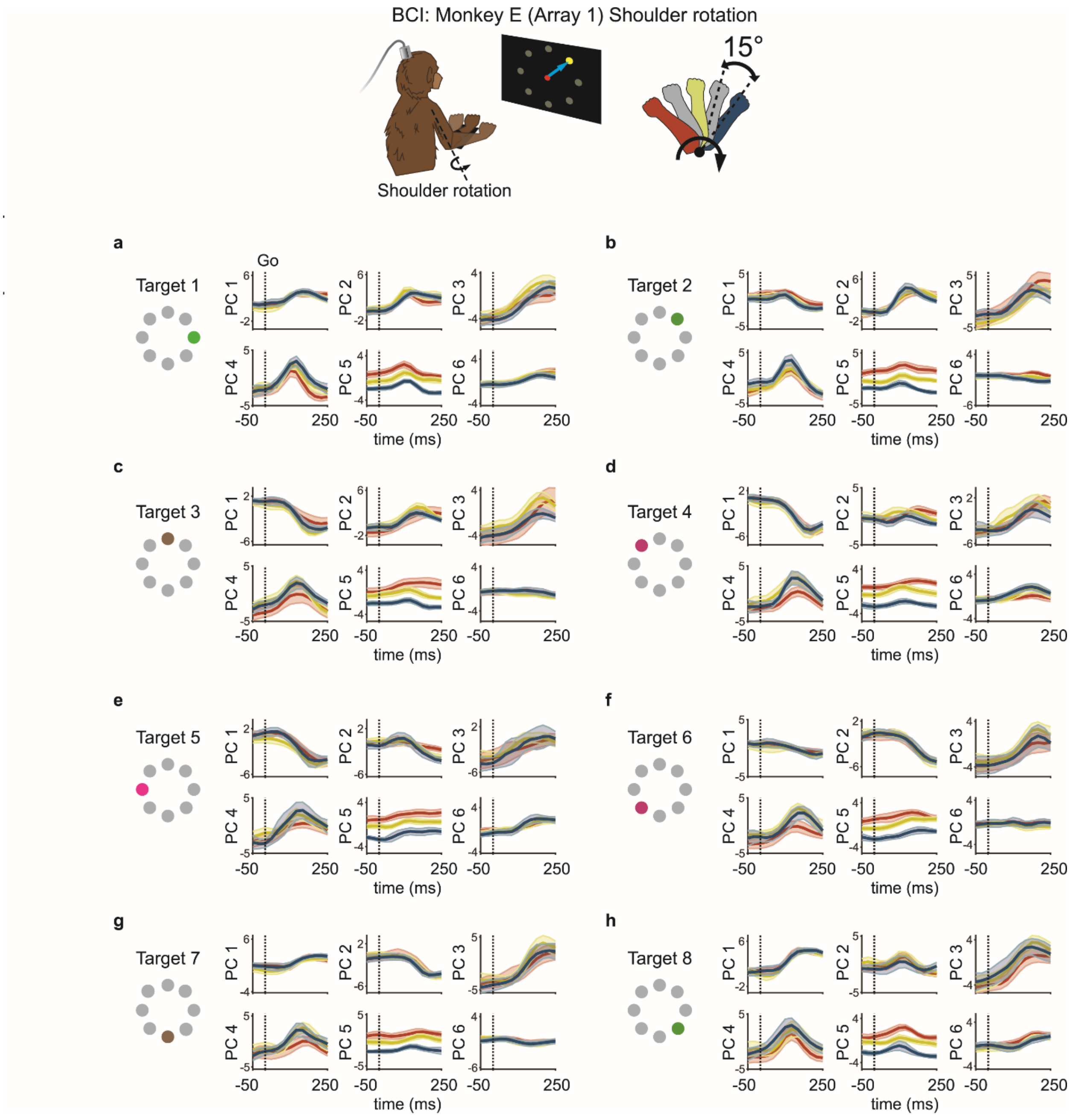


**Supplementary Figure 2 | Neural trajectories from the same target are approximately parallel to each other but offset by posture.**
**(a)** Trial-averaged neural trajectories for the rightward target in Monkey E's BCI task from 50ms before target onset/go cue until 250ms after target onset/go cue. Neural activity is projected onto PCA dimensions identified using neural activity during the analysis window in Figure 2d (i.e., PCs 1-3 are the same as shown in Figure 2d). Data from 3 postures are shown. For all targets, trajectories are approximately parallel but offset by posture (e.g., red trace and blue trace in Figure 2a are approximately parallel). Shaded region around each trajectory indicates SEM. Dotted vertical line indicates target onset/go cue. **(b-h)** Same as (a) for the other target directions.

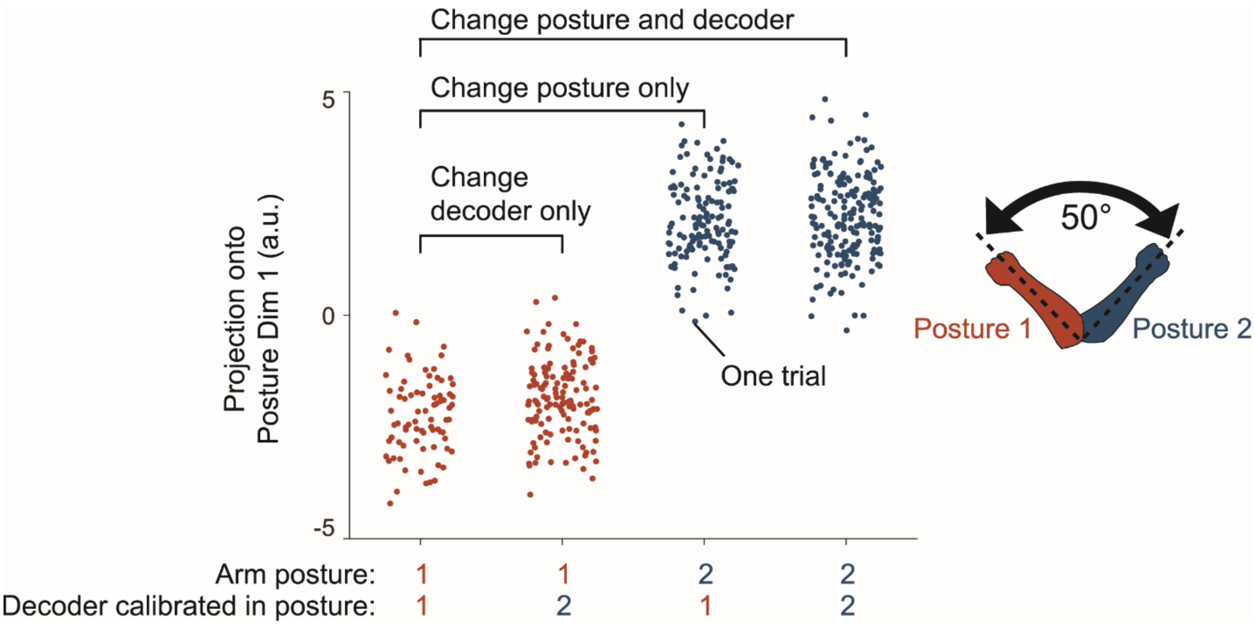


**Supplementary Figure 3 | Postural effects in neural activity are not explained by changes in BCI decoder.**
For the multi-posture BCI paradigm, new decoders were calibrated in each posture. This was done (1) so that proficient control of the BCI could be maintained even if changing arm posture altered neural activity, and (2) so that animals did not attempt to alter their neural activity in one posture to conform to a decoder calibrated in a different posture. However, this experimental design created a potential confound: the observed changes in neural activity across blocks could be due to either changes in arm posture or changes in decoder. To disambiguate between these possibilities, we conducted a control experiment that dissociated the effects of posture and changing decoder. In the experiment, Monkey E performed the center-out BCI task in two postures. Decoders were calibrated in each posture, and the monkey performed the task using each decoder in each posture. Posture was changed by rotating the forearm about the shoulder in the transverse plane by 50 degrees. We refer to the resulting postures as posture 1 and posture 2. We refer to the decoder calibrated in posture 1 as decoder 1, and the decoder calibrated in posture 2 as decoder 2. The monkey then performed four blocks of trials. In block 1, decoder 1 was used in posture 1. In block 2, decoder 2 was used in posture 1. In block 3, decoder 1 was used in posture 2. In block 4, decoder 2 was used in posture 2. We asked whether the changes in neural activity along the putative posture dimension were driven by changes in posture or changes in decoder. To do so, we first used neural activity from blocks 1 and 4 to identify the posture dimension using the same procedure as in Figure 3 (see Methods). Blocks 1 and 4 were chosen because, in these blocks, the animal controlled the cursor using a decoder that was calibrated in the current posture. This matches our procedure for identifying posture dimensions for our main experiments, because in our main experiments, the animal also controlled the cursor using a decoder calibrated in the current posture. We then projected neural activity from each trial (all blocks) onto the posture dimension identified using blocks 1 and 4. For simplicity, the z-scored firing rate estimates for each trial were averaged together over the analysis window to produce a single vector of firing rates (i.e., an $N \times 1$ vector, where $N$ is the number of recorded neural units) for each trial. The analysis window used for each trial was the same as was used for all other BCI tasks (from 50ms after go cue/target onset until 250ms after go cue/target onset). Trials are grouped by experimental block along the horizontal axis. We found changes in neural activity along the posture dimension (vertical axis) result from changing the arm posture, but not from changing the BCI decoder.

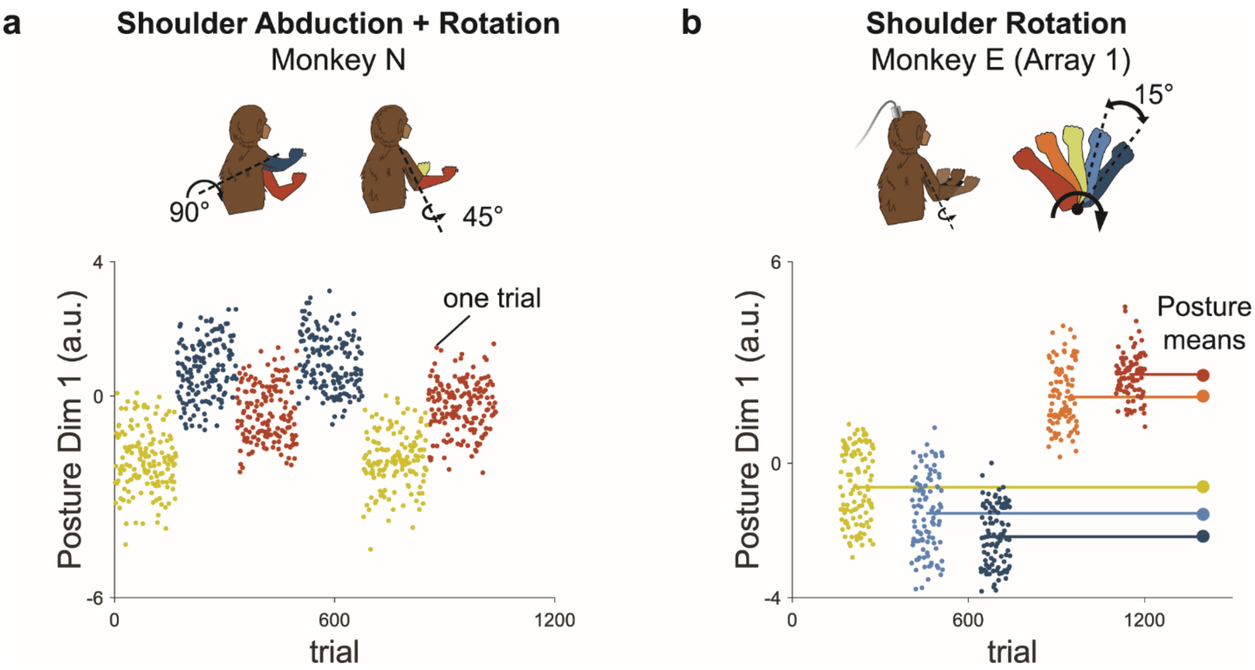


**Supplementary Figure 4 | Postural effects in neural activity are not explained by temporal "drift."** For some multi-posture BCI sessions, the sequence of postures occupied by the arm was ordered by displacement from one extreme. For example, for some shoulder rotation sessions, the arm began in the innermost posture, then was gradually rotated to the outermost throughout the session. This created a potential confound, as putative posture effects in neural activity could have been due to temporal drift in neural activity [S4-S6]. To assess whether changes in neural activity across postures were due to posture or temporal drift, we analyzed data from sessions in which the occupied postures were not ordered. As in Figure 3, we used PCA to identify two posture dimensions. We then plotted the projection of neural activity on the first posture dimension versus trial number. If the changes in neural activity were due to temporal drift, we would expect a steady increase or decrease in the projection along the posture dimension throughout the session. If instead the changes in neural activity were due to posture, the projection along the posture axis would correspond to the posture of the arm, rather than the trial number. **(a)** Projection of neural activity along the posture dimension versus trial number for one of Monkey N's BCI sessions. In this session, shoulder rotation and abduction were used. The projection of neural activity on the posture dimensions changes with arm posture and does not steadily increase or decrease with trial number. This shows that changes observed in neural activity across postures are due to arm posture and not temporal drift. **(b)** Same as (a), but for one of Monkey E's shoulder rotation sessions**.**

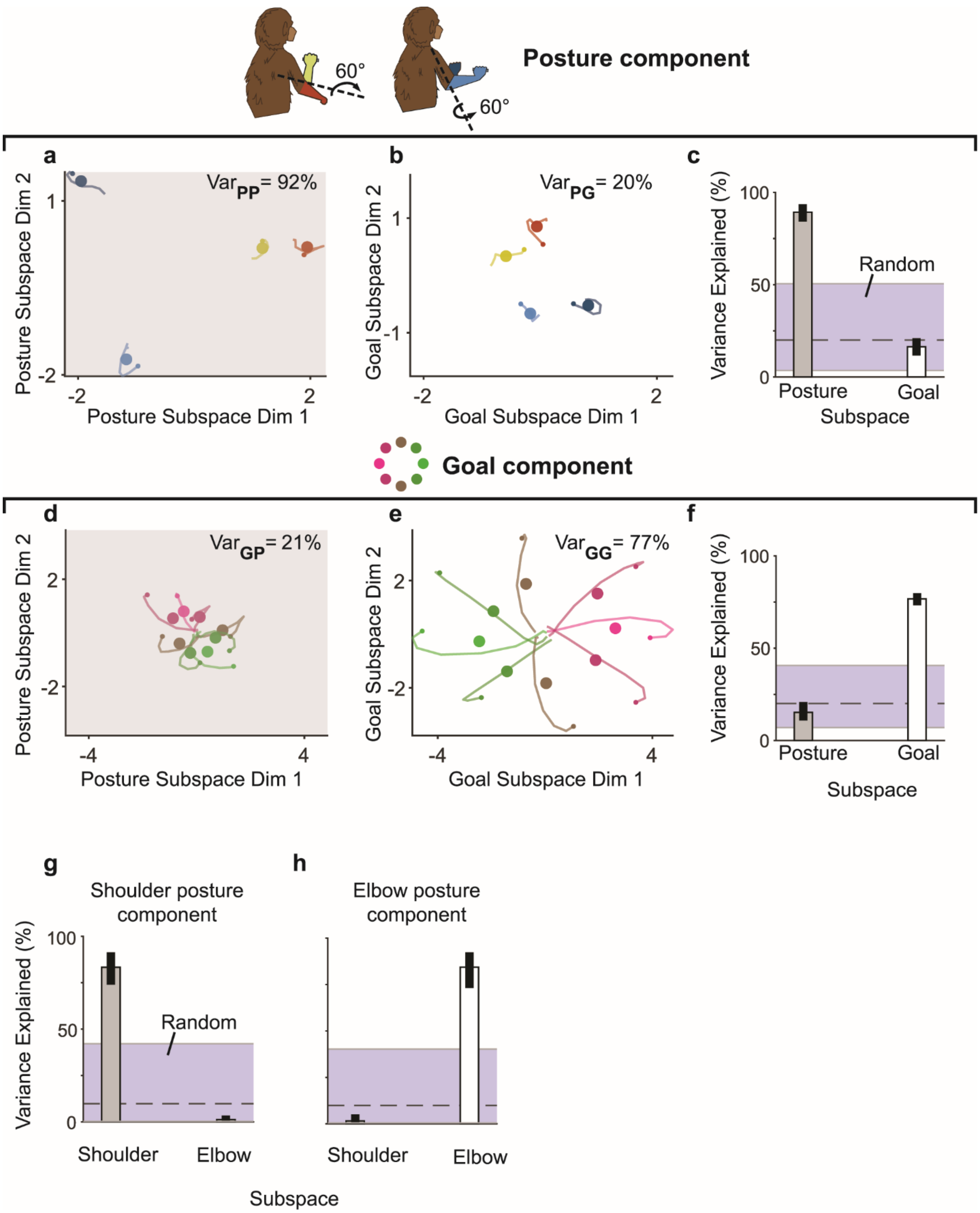


**Supplementary Figure 5 | Elbow and shoulder displacement modulate distinct posture dimensions which are nearly orthogonal to goal dimensions and to each other.**
For all but one of our multi-posture BCI experiments, we applied a postural manipulation of only one joint. We conducted an additional experiment with Monkey E in which we manipulated two joints, the shoulder and elbow, within the same session. All multi-posture BCI experiments, including this 2-joint session, were

included in Figure 3 for completeness. Here, we further analyze the 2-joint Monkey E session. Specifically, we ask how aligned the combined (shoulder and elbow) posture subspace was with the goal subspace, and how aligned the shoulder subspace was with the elbow subspace. **(a)** Projection of the combined posture component of neural population activity onto the posture subspace (same conventions as Figure 3). For this session, the posture subspace captured 92% of the posture-related variance. **(b)** Projection of the combined posture component of neural population activity onto the goal subspace. The goal subspace captured 20% of the posture-related variance. **(c)** Posture-related variance captured by posture and goal subspaces (same format as Figure 3d). The goal subspace captured much less posture-related variance than the posture subspace ($p < 10^{-92}$, bootstrap test), and the amount of posture-related variance captured by the goal subspace was on the low end of the random subspace distribution. **(d-f)** Same format as (**a-c**), but for goal-related variance. The posture subspace captured much less goal-related variance than the goal subspace ($p < 10^{-27}$, bootstrap test), and the amount of goal-related variance captured by the posture subspace was on the low end of the random subspace distribution. Together, (**a-f**) indicate that the combined posture subspace and goal subspace are nearly orthogonal. **(g-h)** We next assessed the alignment between the posture subspaces produced by shoulder and elbow manipulation. To do this, we first split the data into two different groups: trials with different shoulder postures (trials from the dark and light blue postures in the schematic at the top of Figure S5), and trials with different elbow postures (red and yellow postures in the schematic at the top of Figure S5). We identified the ‘shoulder posture component’ by averaging neural trajectories from the shoulder group over different targets. We then identified the ‘shoulder posture subspace’ by applying PCA to the shoulder posture component. We applied an analogous procedure to identify the ‘elbow posture component’ and ‘elbow posture subspace’ from the elbow group. With these components and subspaces identified, we applied the cross-projection alignment test exactly as in Figure 3 to measure the alignment of the shoulder and elbow posture subspaces. The elbow subspace captured much less shoulder-related variance than the shoulder subspace ($p < 10^{-68}$, bootstrap test), and the amount of shoulder-related variance captured by the elbow subspace was on the low end of the random subspace distribution. The shoulder subspace captured much less elbow-related variance than the elbow subspace ($p < 10^{-43}$, bootstrap test), and the amount of shoulder-related variance captured by the elbow subspace was on the low end of the random subspace distribution. Together, these indicate that the shoulder and elbow subspaces are nearly orthogonal.

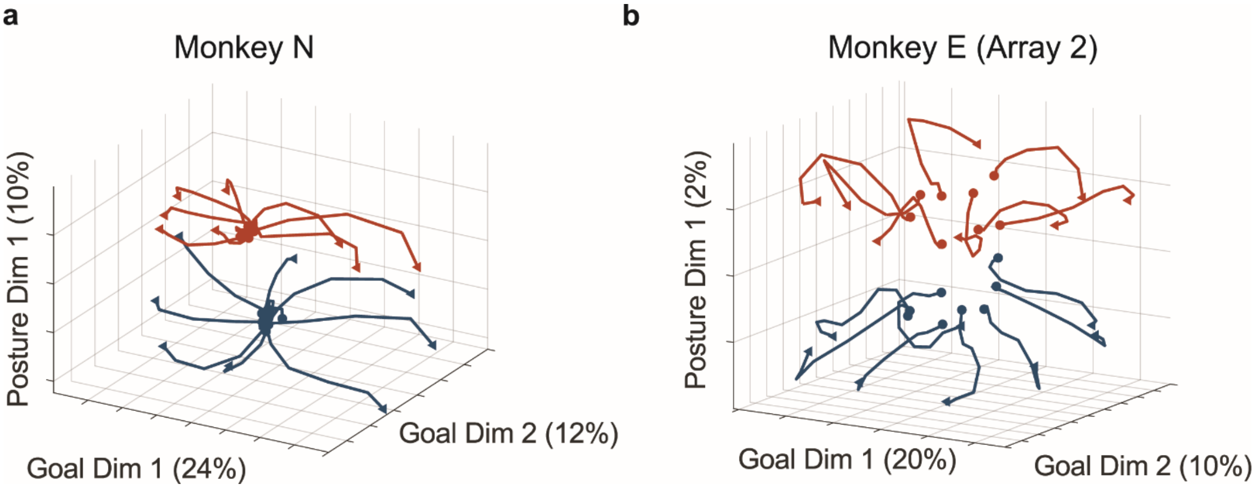


**Supplementary Figure 6 | Neural population trajectories in the reaching task for other monkeys.** Same format as Figure 4e, but for Monkeys N and E. For all monkeys, posture and movement goal modulated separate neural dimensions (cf. Figure 4e).

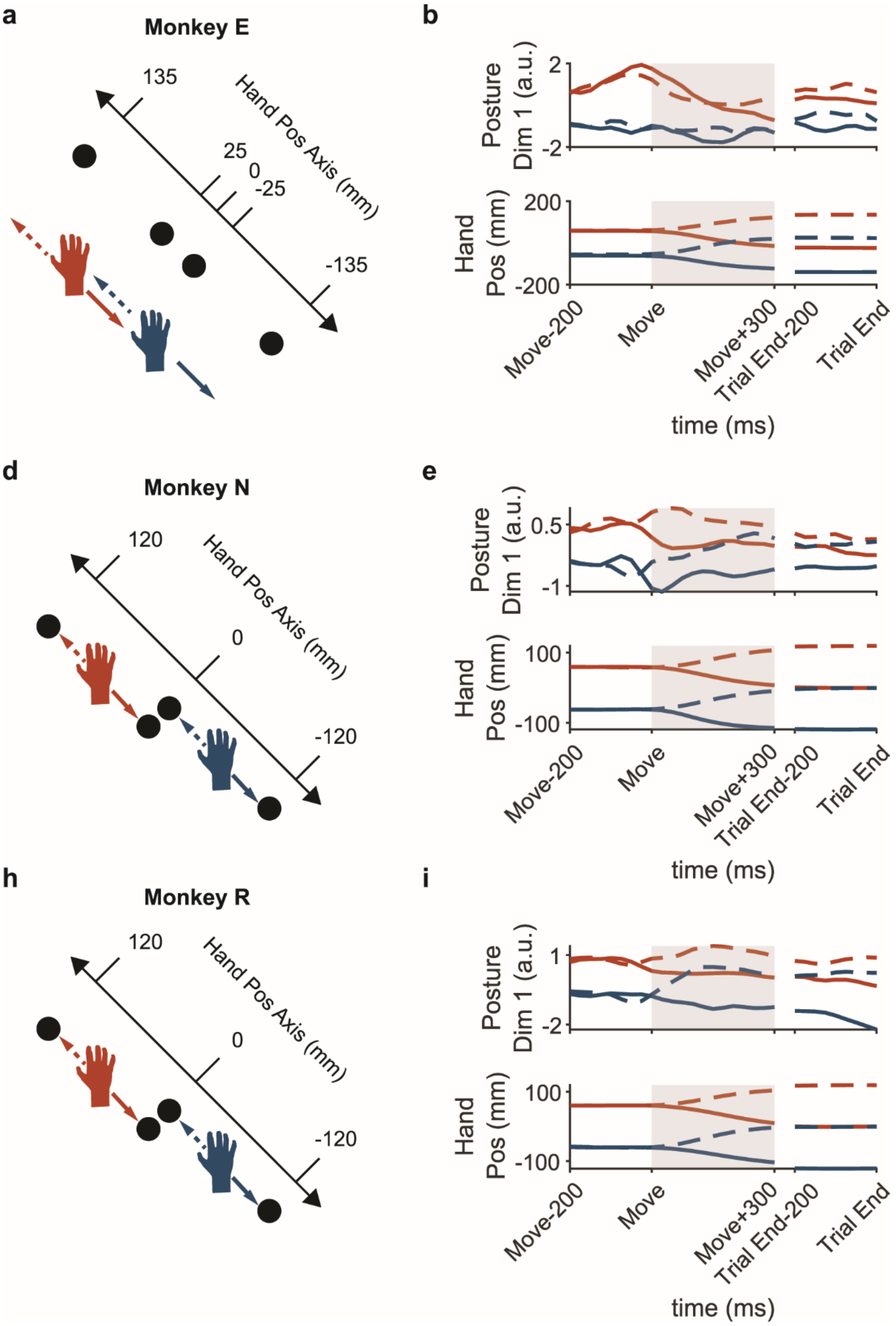

a
Monkey E
135
25
0
-25
-135
Hand Pos Axis (mm)
b
Posture Dim 1 (a.u.)
2
-2
Hand Pos (mm)
200
-200
Move-200
Move
Move+300
Trial End-200
Trial End
time (ms)
d
Monkey N
120
0
-120
Hand Pos Axis (mm)
e
Posture Dim 1 (a.u.)
0.5
-1
Hand Pos (mm)
100
-100
Move-200
Move
Move+300
Trial End-200
Trial End
time (ms)
h
Monkey R
120
0
-120
Hand Pos Axis (mm)
i
Posture Dim 1 (a.u.)
1
-2
Hand Pos (mm)
100
-100
Move-200
Move
Move+300
Trial End-200
Trial End
time (ms)

**Supplementary Figure 7 | Neural activity in the posture subspace does not closely track hand position during movement.**

We asked whether neural activity in the posture subspace tracked hand position during reaching. To do so, we analyzed reaches in which the hand moved along the direction that separated two initial arm postures. We used targeted dimensionality reduction to identify a posture dimension as in Figure 4, then projected trial-averaged neural activity onto this dimension. Because the posture dimension captured changes in neural activity associated with changing arm posture between the two initial locations, we reasoned that neural activity along the posture dimension may continuously track hand position as it moved along the spatial direction separating the two initial arm postures. We refer to the spatial direction separating the two initial arm postures as the “Hand Position Axis.” A position of 0 along the Hand Position Axis corresponds to the location halfway between the two initial postures. **(a)** For Monkey E, we analyzed 4 task conditions: reaching up-left or down-right from the two initial postures shown. Black circles indicate target locations. **(b)** (top) Trial-averaged neural activity for each condition projected onto the posture dimension. (bottom) Trial-averaged hand position for each condition projected onto the hand position axis shown in (a). Neural activity along the posture dimension does not closely track the arm position. **(d-i)** Same as (a-b) for monkeys N and R.

1) Subsample neural trajectories without replacement from each condition into two groups. Take averages for each group.

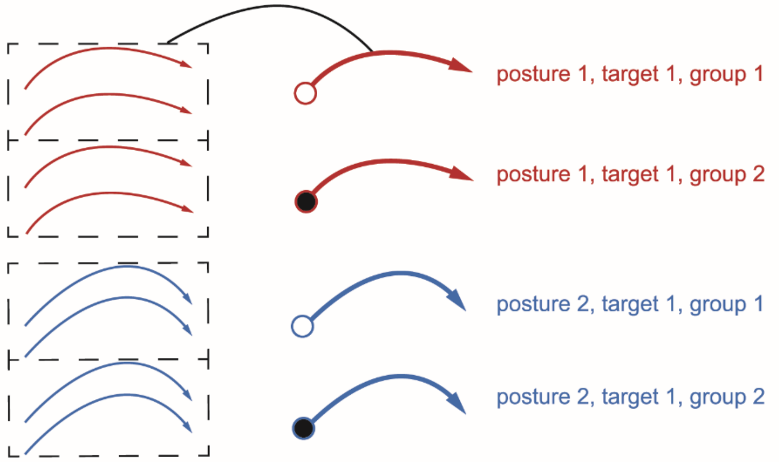


2) Measure across-condition difference before and after applying translation (mean shift)

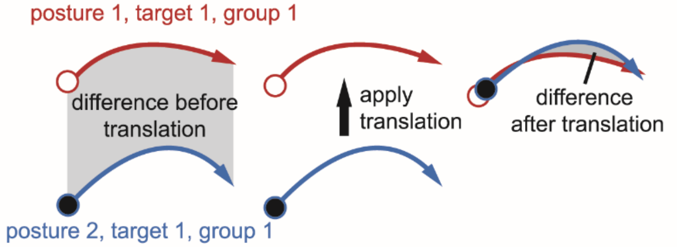


3) Measure within-condition difference

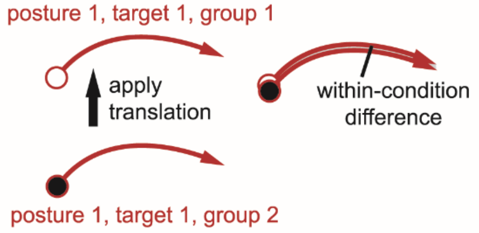


4) Compute normalized difference before and after shift

$$\text{Difference before translation (normalized)} = \frac{\text{Difference before translation}}{\text{Within-condition difference}}$$

$$\text{Difference after translation (normalized)} = \frac{\text{Difference after translation}}{\text{Within-condition difference}}$$

**Supplementary Figure 8 | Method for assessing similarity in neural trajectories across postures.** To assess the similarity of neural trajectories across postures (Figure 6), we asked how well pairs of trajectories from different postures matched if we translated them to bring them into alignment. If a pair matched perfectly after translation, this would indicate that the trajectories had the same shape but started in different locations due to posture-related effects in neural activity. **1)** For each comparison, we first randomly sampled trials from each condition without replacement to form two groups (e.g., condition 1 group 1, condition 1 group 2, condition 2 group 1, condition 2 group 2). We then computed trial-averaged trajectories for each group. Dividing into groups in this way allowed us to measure the difference after translation between trajectories from different groups of the same condition (e.g., condition 1 group 1 and condition 1 group 2), which provided a lower bound for our difference measurements. **2)** We then measured the difference between the condition 1 group 1 mean trajectory and condition 2 group 1 mean trajectory before and after translation. The difference between the pair of trajectories was computed by measuring the Euclidean distance between the trajectories at each timestep and averaging those distances across timesteps. To apply the translation, the trajectory from condition 2 was translated by the vector connecting its mean (taken across timesteps) to the mean (taken across timesteps) of the trajectory from condition 1. **3)** Next, we computed the difference after translation between the trajectories from condition 1 group 1 and condition 1 group 2, which we refer to as the 'within-condition' difference. This measurement provided a lower bound for our other difference measurements. **4)** To combine results across conditions and monkeys, we normalized all differences by the within-condition difference.
